# A Snapshot of the Global Drinking Water Virome: Diversity and Metabolic Potential Vary with Residual Disinfectant Use

**DOI:** 10.1101/2021.10.07.463401

**Authors:** Bridget Hegarty, Zihan Dai, Lutgarde Raskin, Ameet Pinto, Krista Wigginton, Melissa Duhaime

## Abstract

Viruses are important drivers of microbial community ecology and evolution, influencing microbial mortality, metabolism, and horizontal gene transfer. However, the effects of viruses remain largely unknown in many environments, including in drinking water systems. Drinking water metagenomic studies have offered a whole community perspective of bacterial impacts on water quality, but have not yet considered the influences of viruses. In this study, we address this gap by mining viral DNA sequences from publicly available drinking water metagenomes from distribution systems in six countries around the world. These datasets provide a snapshot of the taxonomic diversity and metabolic potential of the global drinking water virome; and provide an opportunity to investigate the effects of geography, climate, and drinking water treatment practices on viral diversity. Both environmental conditions and differences in sample processing were found to influence the viral composition. Using free chlorine as the residual disinfectant was associated with clear differences in viral taxonomic diversity and metabolic potential, with significantly fewer viral populations and less even viral community structures than observed in distribution systems without residual disinfectant. Additionally, drinking water viruses carry antibiotic resistance genes (ARGs), as well as genes to survive oxidative stress and nitrogen limitation. Through this study, we have demonstrated that viral communities are diverse across drinking water systems and vary with the use of residual disinfectant. Our findings offer directions for future research to develop a more robust understanding of how virus-bacteria interactions in drinking water distribution systems affect water quality.

## 1. Introduction

In an effort to prevent the spread of waterborne diseases, previous research on viruses in drinking water has focused on tracking and eliminating human viral pathogens (Dong et al., 2010; Gall et al., 2015; Petrovich et al., 2020; Rao et al., 1981; Ye et al., 2012). For example, researchers have tracked specific human viruses with qPCR to understand their distribution throughout drinking water treatment plants and distribution systems (Albinana-Gimenez et al., 2006; Samhan et al., 2015; Ye et al., 2012). While research on the presence and fate of human viruses is essential for assessing drinking water quality, the impacts of bacterial viruses, commonly referred to as bacteriophages (or phages), have rarely been addressed. Phages influence microbial populations (Rohwer et al., 2009; Suttle, 2007) and biogeochemical cycling (Parmar et al., 2017), through mortality (Fuhrman and Noble, 1995; Guidi et al., 2016) and by facilitating horizontal gene transfer between microorganisms and across environments (Rohwer et al., 2009). Similarly, viruses likely impact both beneficial and detrimental bacteria and other microorganisms in drinking water systems that have both direct (e.g., causing illness) and indirect effects (e.g., nitrification and corrosion) on drinking water quality.

Metagenomic studies have become critical to evaluate the cumulative effects of the drinking water microbiome on water quality (Hull et al., 2019; Liu et al., 2013). Past studies have shown that many factors, ranging from source waters (Assche et al., 2019; Stanish et al., 2016) to treatment practices (Bautista-de los Santos et al., 2016; Dai et al., 2020; Pinto et al., 2012), influence the bacterial, archaeal, and eukaryotic inhabitants of drinking water. For instance, the use of residual disinfectants (e.g., free chlorine, chloramine) has been connected to the selection of opportunistic pathogens (Feazel et al., 2009; Kotlarz et al., 2019; Liu et al., 2019; Vaerewijck et al., 2005), the prevalence of antimicrobial resistance genes (Jia et al., 2019, 2015; Shi et al., 2012), and shaping microbial communities (Dai et al., 2020; Hull et al., 2019; Krishna et al., 2020; Wang et al., 2014). At most, these studies have quantified the fraction of viral sequences in a cellular metagenome (Brumfield et al., 2020; Gomez-Alvarez et al., 2012; Stamps et al., 2018). This leaves fundamental information about drinking water viruses, such as their taxonomic or metabolic diversity and influence on the drinking water microbiome, unknown. To fill similar knowledge gaps in other environments, cellular metagenomes have been re-examined for DNA viruses, revealing novel viral diversity and offering insights into viral-host interactions (Paez-Espino et al., 2016; Roux et al., 2015b). Taking such an approach can provide an untapped opportunity to illuminate the effects of phages on drinking water quality and safety.

The overall goal of this work was to characterize viral communities in drinking water distribution systems. To do so, we identified viral sequences from publicly available metagenomic datasets from drinking water distribution systems from around the world to assess viral taxonomic diversity and metabolic potential. We further investigated the effect of study characteristics (e.g., the type of filter used for sample processing and the DNA extraction method) and environmental conditions (e.g., water quality) on the structuring of the observed viral communities. This study lays the groundwork for understanding the impact of viruses on the microbial communities in drinking water distribution systems.

## 2. Methodology

### 2.1 Data Collection

Publicly available drinking water metagenomic reads were downloaded from NCBI using the SRA Toolkit v2.9.6 (Leinonen et al., 2011) (Table S1). Sequences deposited on MG-RAST were directly requested from researchers. Only water samples that were collected from drinking water distribution systems and that were sequenced using Illumina short read technology were included in our analysis, excluding samples taken from within treatment plants and distribution system biofilm samples. Based on these criteria, 164 samples were used. Sequencing for some samples was replicated, resulting in a total of 312 sequencing runs. Replicate sequencing sets from the same samples were pooled during subsequent analyses (“sample” Table S1). Table S1 provides information about sample processing, sequencing platform, and other metadata for each sample included in this study. Water quality parameters for the Dai et al (2020) and Sevillano et al (2021) samples are provided in Tables S2 and S3, respectively.

### 2.2 Sequence Read Processing

Sequence reads were filtered and trimmed using fastp v0.20.1 (Chen et al., 2018). All code is deposited on github and made freely available (https://github.com/DuhaimeLab/global_drinking_water_viromes_Hegarty_et_al_2021). The UniVec Core database (as of February 2021) was used to remove contamination from vectors (NCBI, 2016).

### 2.3 Assembly

The filtered and processed reads from each distribution system sampling location were co-assembled using MetaSPAdes v3.10.1 (Nurk et al., 2017). Assembly quality was assessed with QUAST v5.0.2 (Gurevich et al., 2013).

### 2.4 Viral Population Identification

To define viral populations, putative viral contigs were identified from contigs greater than 3 kb using a strategy based on VirSorter v1.0.6 (Roux et al., 2015a), VirSorter2 v2.1 (Guo et al., 2021), CheckV v0.7.0 (Nayfach et al., 2020), VIBRANT v1.2.1 (Kieft et al., 2020), and VirFinder v1.1 (Ren et al., 2018) predictions (Figure S2). Using the predictions considered high quality by the tools’ authors, any contig classified (i) as category 1 or 2 by VirSorter, (ii) as high or medium by VIBRANT, (iii) as complete, high, or medium by CheckV, (iv) with a VirSorter2 score greater than 0.95, (v) with at least two hallmark viral genes from VirSorter2, or (vi) with a VirFinder score greater than 0.9 were considered viral. Additionally, any contig called viral by at least two of the following were considered viral: VirSorter categories 3-6, VIBRANT low, CheckV low, Virfinder score between 0.7 and 0.9, and VirSorter2 score between 0.5 and 0.95. If a contig had (i) no viral genes and more than one host gene according to CheckV, (ii) more than 3 times as many host genes as viral genes based on CheckV, or (iii) a length greater than 50 kb and no viral hallmark genes from VirSorter2, it was removed from the viral list. Bacterial genes were excised from the edges of proviruses based on their identification by CheckV, VirSorter2, and VIBRANT. Based on the community-established standard for defining viral populations (Roux et al., 2019), all viral contigs were clustered (stampede-clustergenomes) (Roux and Bolduc, 2017) if they shared average nucleotide identity (ANI) of 95% across 85% of the contig length (“viral populations” in the following).

The annotations of the three most ubiquitous VPs (based on frequency of occurrence and relative abundance) were examined to assess the confidence of the phage annotations of each of these. These three ubiquitous VPs were blasted against the nucleotide collection (nt) in NCBI to assess whether they matched any previously sequenced proviruses. The top hits were run through CheckV to identify any provirus sequences in their genome.

### 2.5 Read Mapping

Filtered and trimmed reads were mapped to all contigs from that sample’s assembly using Bowtie2 (Langmead and Salzberg, 2012) and quantified using SAM Tools v1.11 (Li et al., 2009). To quantify the reads mapped to the trimmed viral sequences, BLAST v2.9.0 (NCBI, 2019) was used to align the trimmed viral sequences to the contigs and then reads overlapping this region were quantified using FeatureCounts (Liao et al., 2014) from the Subread package (Liao et al., 2013).When more than one sequencing run was performed for a given sample, the total read coverage was summed across sampling replicates.

### 2.6 Alpha and Beta Diversity

Only contigs with reads covering at least 3 kb of their length were included in diversity analyses. Alpha diversity measures (i.e., richness and Shannon Dissimilarity) were calculated using the vegan v2.5-7 package (Oksanen et al., 2020) in R v4.0.2 (R Core Team, 2021). All analyses were performed using RStudio v1.3. 595 (R Studio Team, 2020). To normalize for differences in sequencing depth, taxa per one hundred thousand viral reads (henceforth, “normalized richness”) and the Shannon Evenness Index (calculated by dividing the Shannon Dissimilarity by the maximum possible Shannon Dissimilarity for that sample) were used to examine alpha diversity trends.

Transcripts per kilobase million (TPM) was determined for each viral population and used to calculate the Bray Curtis distance between samples using the vegandist function. Permutational Multivariate Analysis of Variance (PERMANOVA) using the adonis function (vegan) was used to test the effect of processing procedures, geography, and residual disinfectant use on the full community structure (Bray Curtis dissimilarities ∼ continent * Köppen zone * filter pore size for biomass collection * DNA extraction method; with stratas for ‘study’). The effect of ‘study’ and residual disinfectant were also tested in isolation (Bray Curtis dissimilarities ∼ study; Bray Curtis dissimilarities ∼ residual disinfectant).

To avoid the confounding effects of study design and climate on diversity analyses, the Dai et al. (2020) samples collected in the temperate oceanic climate zone (Cfb Köppen climate classification scheme) were used to evaluate the effects of residual disinfectant use on taxonomic diversity and metabolic potential (Figure S1 inset). This study was chosen because it provided detailed water quality parameters and was the only study to include samples from distribution systems operated without a residual disinfectant. The six samples taken from the warm summer continental zone (Dfb Köppen climate classification) were excluded because they were all taken from distribution systems that use a residual disinfectant, complicating the isolation of climate’s effect from that of residual disinfectant. Additionally, only samples that used free chlorine (and not chloramines) were included to facilitate comparison with the results presented by Dai et al. (2020). This reduced set of samples was used for all subsequent comparisons of water quality and residual disinfectant. Differences in alpha diversity between distribution systems based on residual disinfectant use were tested using the Wilcoxon rank sum test.

Additionally, both the Sevillano et al. (2021) and Dai et al (2020) samples were used to investigate the effects of water quality parameters, including residual disinfectant, on the viral community. These two studies were chosen as the only two to include both samples with and without residual disinfectant. Distance-based Redundancy Analysis (db-RDA) was performed using the capscale function in the vegan package v2.5-7 (Oksanen et al., 2020) in R (Bray Curtis dissimilarities ∼ chlorine + phosphate + temperature + pH + conductivity + ammonia + nitrate + DO + TOC). PERMANOVA was used to test the effect of these water quality parameters and the drinking water distribution system on the viral community structures of the Dai et al. (2020) samples (Bray Curtis dissimilarities ∼ chlorine + phosphate + temperature + pH + conductivity + ammonia + nitrate + dissolved oxygen (DO) + total organic carbon (TOC) + drinking water distribution system location). Parameters were considered significantly correlated if p-value ≤ 0.05.

### 2.7 Metabolic Potential Analyses

Viral contig gene metabolic annotations were assigned using DRAM based on the KEGG and Pfam databases (Shaffer et al., 2020). Read coverages of the open reading frames predicted by DRAM were calculated using FeatureCounts (Liao et al., 2014) from the Subread package (Liao et al., 2013). Bray Curtis distances between samples were calculated using the vegdist function and then db-RDA was performed using the capscale function with the vegan package in R (Bray Curtis dissimilarities ∼ chlorine + phosphate + temperature + pH + conductivity + ammonia + nitrate + DO + TOC). The water quality parameters were used in a PERMANOVA model to evaluate the effect of water quality parameters on the abundance of KEGG orthologies (KOs), which are molecular function classes that represent the metabolic potential of the viromes. The following equation was used in the adonis function (vegan): Bray Curtis dissimilarities ∼ chlorine + phosphate + temperature + pH + conductivity + ammonia + nitrate + DO + TOC + drinking water distribution system location). Parameters were considered significantly correlated if p-value ≤ 0.05. KEGG pathway modules were considered more abundant in a condition if the difference in normalized read counts was statistically significant between the two conditions (Welch’s Two Sample t-test p-value ≤ 0.05).

### 2.8 Taxonomy

The open reading frames (ORFs) from DRAM were clustered with all full-length bacterial and archaeal virus genomes from NCBI’s viral reference dataset (‘Viral RefSeq’, v.85, January 2018) using vConTACT2 (Bin Jang et al., 2019). Clusters with at least one taxonomic assignment from vConTACT2, as well as clusters found in ≥ 10% of samples, were separately visualized with Cytoscape v3.8.2 (Shannon et al., 2003). The lowest common ancestor was identified for each vContact2 cluster with a taxonomic assignment.

## 3 Results/Discussion

To produce a snapshot of the virome in drinking water distribution systems, we mined publicly available drinking water metagenomes (Chao et al., 2013; Dai et al., 2020; Douterelo et al., 2018; Garner et al., 2018; Jia et al., 2015; Ma et al., 2019, 2017; Potgieter et al., 2020; Sevillano et al., 2021; Shi et al., 2012; Zhang et al., 2019). Table 1 summarizes the metadata for each study. The majority of the samples were collected in Europe (31%) and Asia (35%), with the remainder obtained from North America (23%) and Africa (11%) (Figure S1). In total, 164 samples were collected from 69 distribution systems, with 18 samples from systems with no residual disinfectant and 102 from systems with a residual disinfectant. 46 samples are presumed to be from distribution systems using a residual disinfectant based on the typical disinfection choice of their country of origin (i.e., China, Singapore, South Africa, United Kingdom, United States), but this could not be confirmed (Table S1).

**Table 1.**
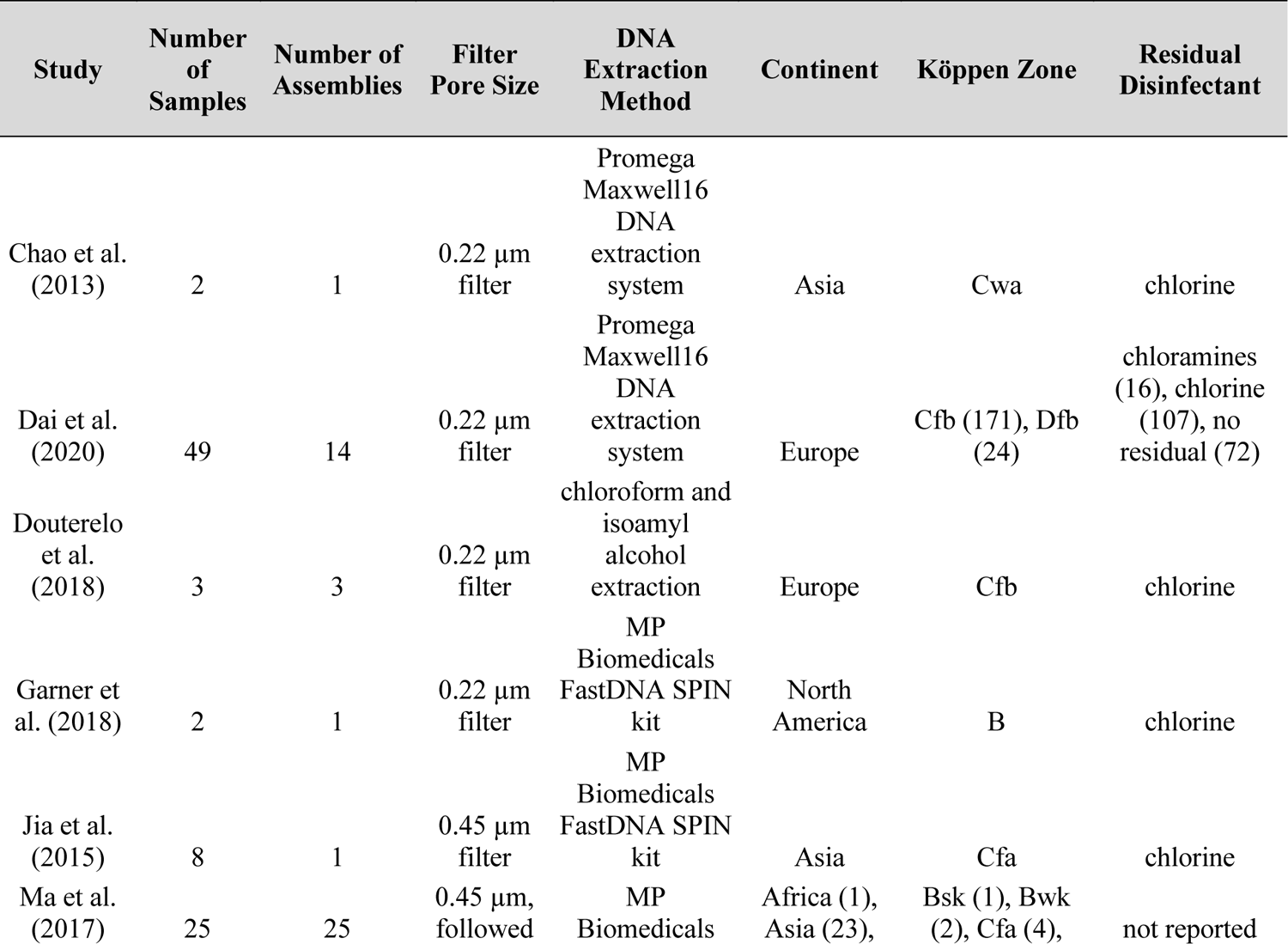
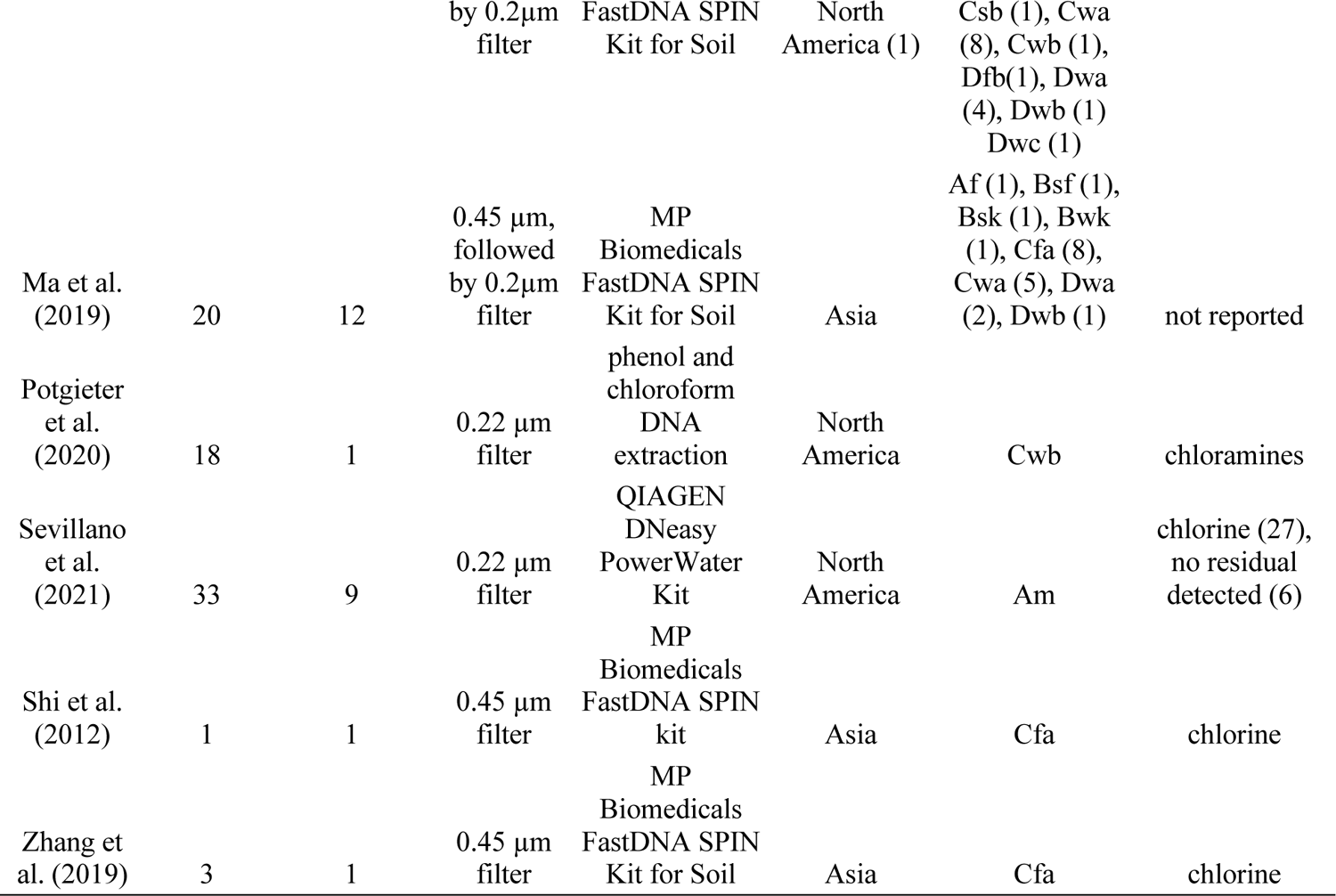
Summary data for the assemblies organized by study.

All samples from the same drinking water distribution system were co-assembled (Tables S1 and S4), resulting in 729,724 contigs larger than 3 kb. We then mined the assembled contigs for viral sequences, identifying 62,500 viral sequences, or 9% of all the contigs longer than 3 kb. The proportion of viral contigs is typically not reported in studies that have mined metagenomes for viral sequences. Our results are in agreement with the upper range (0.3-16%) of the percentage of viral sequences observed in a marine metagenome study (Delong et al., 2006). Additionally, this is consistent with our expectations that viral genomic material would represent roughly 10% of community DNA given that viruses are on average an order of magnitude more abundant than their microbial hosts in most aquatic habitats, while their genomes tend to be one to two orders of magnitude smaller. To facilitate comparisons of the viral populations across distribution systems, the viral contigs were clustered into 48,935 viral populations based on 95% sequence similarity over 85% of the contig length (Roux et al., 2019). These viral populations allowed us to characterize the taxonomic diversity and metabolic potential of the drinking water virome, as well as the geographic and treatment-based factors shaping these communities.

### 3.1 Viral ecology of drinking water communities

#### 3.1.1 Drinking water viral populations are predominantly novel and heterogeneous between samples and studies

We used vConTACT2 to examine the evolutionary relatedness of the viral populations (VPs). Of the total 62,500 viral sequences, 23,573 (37%) formed 7,897 clusters that approximate genus-level relatedness (Bin Jang et al., 2019) (Figure 1A; Figure S3). Only 778 viral clusters (10% of all viral clusters) were given a definitive taxonomic assignment per the International Committee on Taxonomy of Viruses (ICTV) conventions by vConTACT2 (Figure S4). The vast majority of these, 763 out of the 778 viral clusters, were assigned to families in the Caudovirales order of tailed dsDNA phage. *Podoviridae* (373 VPs), *Siphoviridae* (284), and *Myoviridae* (81), were the three most prevalent families of viruses identified in these samples, mirroring the three dominant families in many other freshwater environments (Moon et al., 2020; Saparbaevna et al., 2017; Skvortsov et al., 2016; Zaouri et al., 2020). The dominance of Caudovirales among the classified VPs in our study may represent compositional bias towards this viral order, as the vConTACT reference database is 90% Caudovirales. However, the family classification of our VPs does not reflect database bias, as vConTACT contains 50% *Siphoviridae*, 23% *Myoviridae*, and 15% *Podoviradae* (Bin Jang et al., 2019).

**Figure 1:**
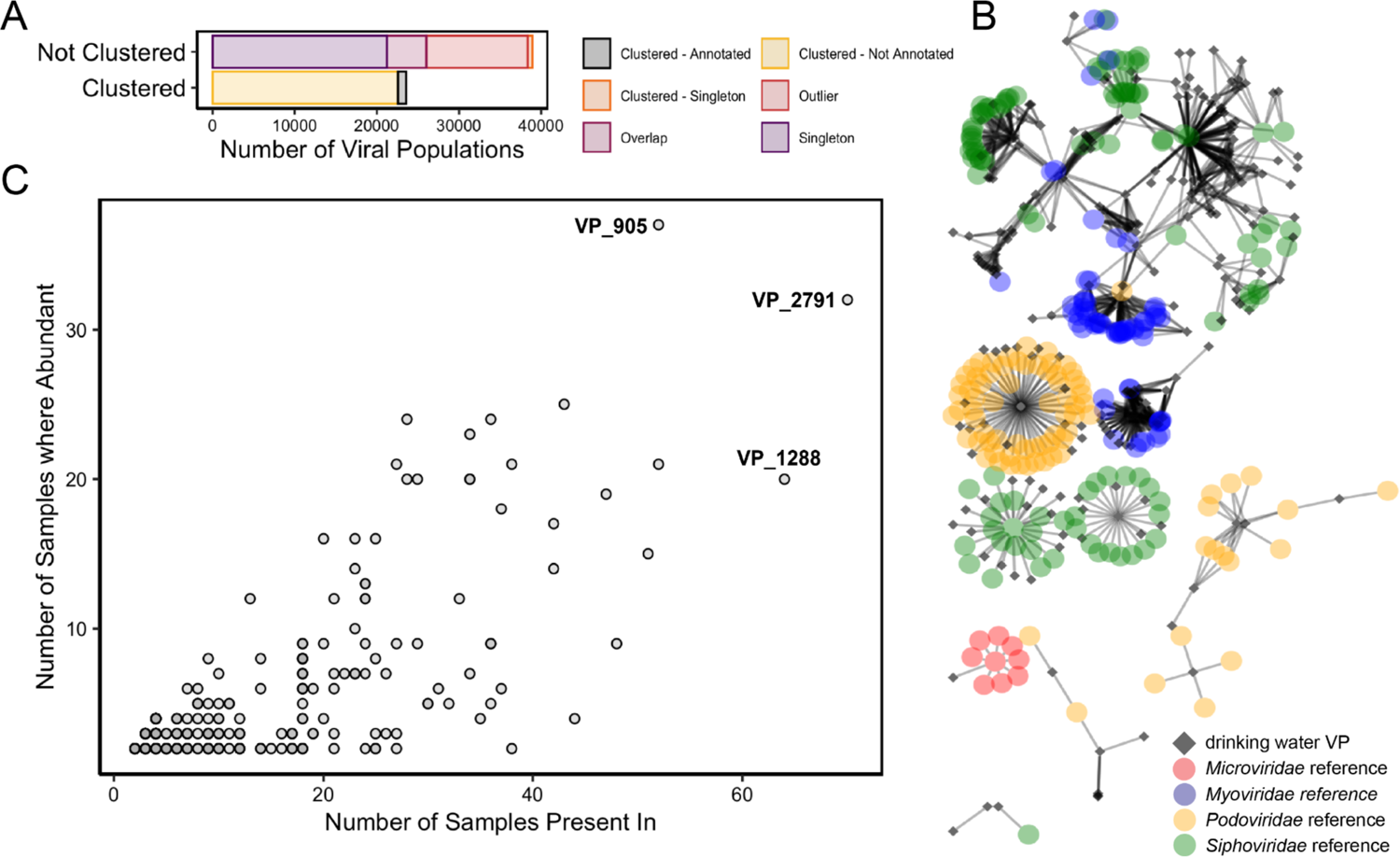
Overview of the drinking water viral populations (VPs). (A) Status of drinking water VPs after taxonomic clustering by vConTACT2. VPs that were given a taxonomic assignment by vConTACT2 are in black. (B) Network clusters of VPs found in more than 10% of samples and their connections to viral reference genomes from vConTACT2. Drinking water VPs are black diamonds and viral reference genomes are colored by family-level taxonomic assignment. (C) Number of samples where a VP was abundant (i.e., greater than 1% of reads) versus the number of samples in which that viral population was found. Only contigs with a relative abundance of greater than 1% in at least one sample were included in this figure. The three most ubiquitous drinking water viral populations were labeled and further described below. Where multiple points overlap, they have a darker fill.

To date, the vast majority of VPs described in environmental viromics studies have been novel, reflecting that viral databases insufficiently represent global viral diversity, as well as the fact that the viral genomes reconstructed from these datasets tend to be incomplete and thus difficult to classify. For instance, in recent studies of the Arctic Ocean (Gregory et al., 2019), honey bees (Deboutte et al., 2020), and New York City wastewater (Gulino et al., 2020), only 9.8%, 9.5%, and 4% of their viral populations, respectively, received a taxonomic assignment. Only 1% of the drinking water viral populations in our study had a taxonomic assignment (Fig. 2A). Since many viruses are strongly habitat-specific (Paez-Espino et al., 2016; Roux et al., 2012), the likelihood of sequencing novel viruses from an environment poorly represented in the databases is high. In addition to the lack of drinking water virome studies to date, the typical source waters (i.e., groundwater and surface water) for drinking water production have been less frequently studied than marine aquatic environments (Kallies et al., 2019; Kothari et al., 2021; Malki et al., 2020; Moon et al., 2020; Paez-Espino et al., 2016; Roux et al., 2012; Skvortsov et al., 2016).

**Figure 2:**
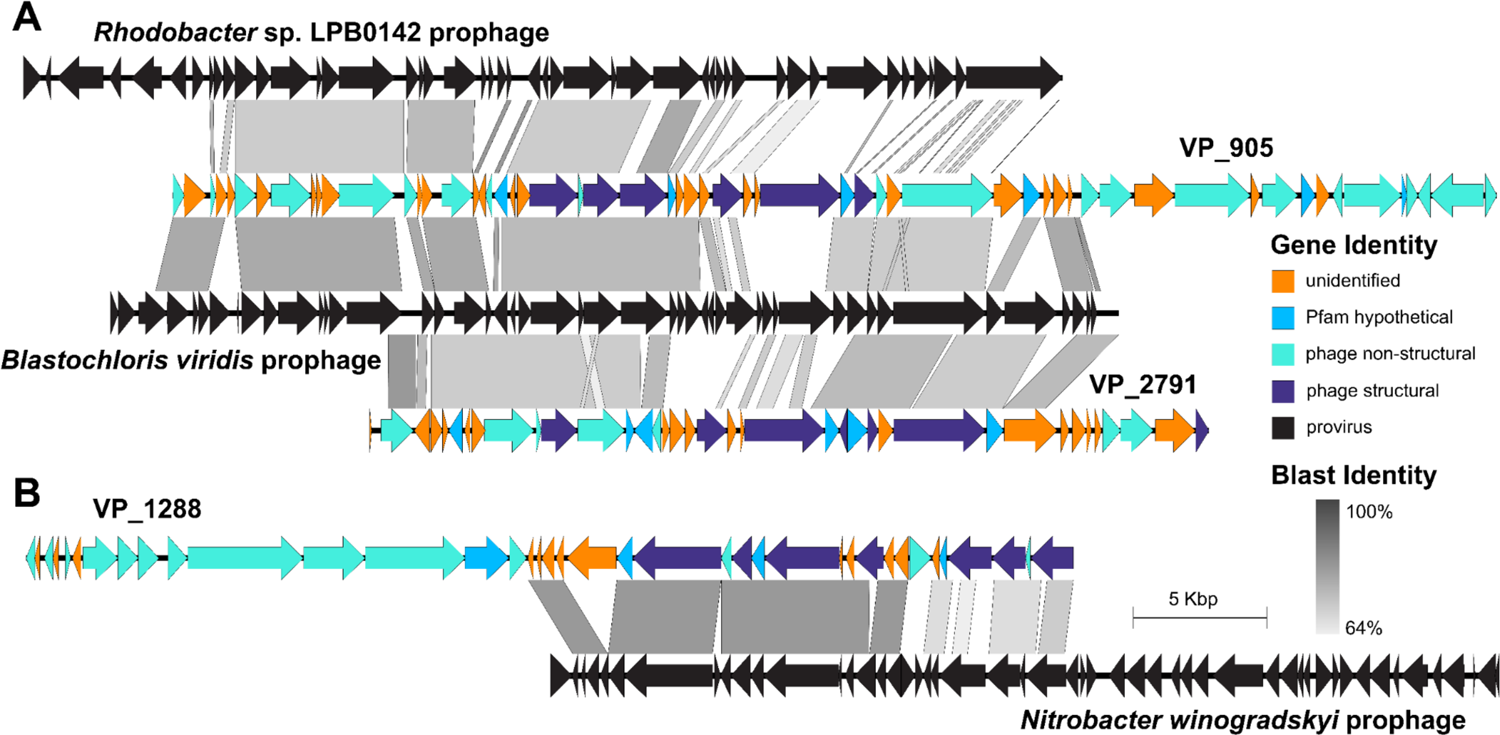
Genome comparisons of the three most globally ubiquitous drinking water viral populations (VPs) with the known sequences (NCBI nt) that share the greatest percent coverage (blastn). The longest representative contig was visualized for each VP. (A) Comparison of VP_905 (55.2 kb) with Rhodobacter sp. LPB0142, comparison of VP 905 and VP_2971 (35.0 kb) with Blastochloris viridis. (B) Comparison of VP_1288 (46.8 kb) with Nitrobacter winogradskyi. Genes on the VPs are colored based on their Pfam identity.

Of the 48,935 VPs (95% ANI across 85% of the contig) identified across all distribution systems, 156 viral populations were found in more than 10% of the samples. Eight of these had an order-level taxonomic assignment: seven belonged to the Caudovirales order (two viral populations of *Myoviridae*, one *Podoviridae*, two *Siphoviridae*, two unassigned families), and one VP belonged to the *Microviridae* family of isometric ssDNA phage in the *Petitvirales* order (Figure 1B). Other studies have similarly found a high degree of heterogeneity with few ubiquitous viral populations across different viromes collected from similar environments (e.g., within ecological zones in marine environments) (Gregory et al., 2019; Paez-Espino et al., 2016; Roux et al., 2016).

The three most ubiquitous viral populations were VP_905, VP_2791, and VP_1288 (Figure 1C). VP_2791, the most prevalent, was found in 42% of the samples. VP_905 and VP_2791 were classified as unknown genera within the *Siphoviridae* family (Caudovirales). VP_1288 did not have a taxonomic assignment at any level. The top blast hits for each of the most ubiquitous VPs in the NCBI nt database (Table S5) were each within prophages identified by CheckV. Both VP_905 and VP_2971 were most similar (based on query coverage) to a sequence in *Blastochloris viridis* (Figure 2A, Table S5). *B. viridis* is a purple photosynthetic bacterium (Tsukatani et al., 2015) that was isolated from a German river and has been found in other freshwaters, including in the biofilm of an ultrapure supply water plant (Bohus et al., 2010).

VP_905 also matched with a similar query coverage to *Rhodobacter* sp. LPB0142 (Figure 2A) and *Paracoccus contaminans.* VP_1288 was most similar to *Nitrobacter winogradskyi*, a nitrite oxidizing bacterium that has been found in soils, as well as freshwater (Mellbye et al., 2015) (Figure 2B). While most of the ORFs on the three contigs were not assigned a metabolic annotation (Figure 2; Table S6), the high number of unknown and phage-related genes, combined with the high sequence similarity and synteny with known prophages, increased the confidence that each are truly phage contigs. By pairing deeply-sequenced viromes with cellular metagenomes, subsequent studies will build on this first report of ubiquitous viral populations in drinking water to establish “who infects whom”, thereby leading to a more robust understanding of how viruses impact drinking water quality.

#### 3.1.2 Environmental conditions and experimental design influence drinking water viral communities

To evaluate the possible effects of experimental design and environmental conditions on viral community structure, we tested the effects of the continent and Köppen climate zone where samples were collected, the filter pore size for biomass collection, and the DNA extraction method. In contrast to differences reported between viruses sequenced from 0.1-20 µm and <0.1 µm fractions (Williamson et al., 2012), our meta-analysis found that filter pore size did not impact viral community structures (PERMANOVA; Table S7). As virus sizes typically range from 0.02-0.3 µm, the relatively large filter pore size (0.22 µm and 0.45 µm) used across all studies in our analysis may explain why filter pore size did not affect viral community structure.

The other three parameters tested, as well as the interaction of the DNA extraction method and Köppen climate zone, did have a statistically significant effect on the viral community’s structure (PERMANOVA; Table S7). Combined, the parameters and their interactions tested explained slightly over a quarter of the variability in the system (residual PERMANOVA R^2^=0.73; Table S7). The inability to explain the full variability in beta diversity in the PERMANOVA model could reflect the unbalanced nature of the metadata (e.g., many more samples used the Maxwell 16 DNA extraction system than the other three methods; Figure S4), as well as the importance of additional factors.

We additionally tested the effect of ‘study’ to encompass the experimental design and environmental conditions, including those not explicitly reported in all of the original metagenomics studies (e.g., water source, distribution system water qualities, library preparation). ‘Study’ was found to significantly impact the viral community structure (PERMANOVA R^2^=0.21, p-value=0.0001). In a previous meta-analysis of drinking water bacterial communities, ‘study’-specific effects explained nearly a third of the variability among samples (Bautista-de los Santos et al., 2016). The collection and processing of samples are known to have a substantial impact on the viruses identified (Brinkman et al., 2018; Dias et al., 2020; Hurwitz et al., 2013; Percival and Wyn-Jones, 2013). The low biomass of drinking water samples may make them particularly prone to the influence of ‘study’-specific differences from contamination during sample processing (Bautista-de los Santos et al., 2016). Alternatively, as most studies only examined one distribution system or focused on a single region, the strong ‘study’-specific effects may be partly due to local factors, such as geography, climate, treatment regime, and water source (e.g., groundwater, river water). Such parameters have been shown to influence the viral communities of many other environments (Paez-Espino et al., 2016).

To investigate the influence of water quality parameters independently from study-specific effects, we focused on the Dai et al. (2020) samples, collected from the UK and Netherlands, and the Sevillano et al. (2021) samples, collected from Puerto Rico. Both studies found that chlorine was a major determinant of the differences in the microbial community composition (Dai et al., 2020; Sevillano et al., 2021). Dai et al. (2020) additionally found that DO and conductivity were correlated with the microbial community structure of these samples; Sevillano et al. (2021) found that DO, temperature, and TOC were each significantly associated with changes in microbial composition at specific sites. As bacterial community structure is a critical driver of bacteriophage community structure (Gregory et al., 2019; Jonge et al., 2018), we hypothesized that these parameters would also be correlated with the viral community relatedness (Bray Curtis dissimilarities). We found that viral beta diversity (i.e., differences in community structure between samples) correlated with chlorine, phosphate, and nitrate concentration, DO, conductivity, water temperature, pH, and TOC for both studies; ammonia concentration was also significantly correlated for the Dai et al. (2020) samples (Figure 3; Tables S8 and S9). These trends indicated that water quality parameters do likely shape the viral communities of drinking water. Distribution system sampling location was significantly correlated with the viral community structure of both the Dai et al. (2020) study (PERMANOVA R^2^=0.33, p-value=0.001; Table S8) and Sevillano et al. (2021) study (PERMANOVA R^2^=0.32, p-value=0.0001; Table S9). Together with the unexplained variability in both the Dai et al. (2020) (PERMANOVA R^2^=0.19; Table S8) and the Sevillano et al. (2021) samples (PERMANOVA R^2^=0.25; Table S9), these trends indicated that unmeasured factors also likely affected the viral community structures. Altogether, the Sevillano et al. (2021) and Dai et al. (2020) viral sequences suggest that water quality parameters are influencing the structure of viral communities independently of study-specific differences in experimental design.

**Figure 3:**
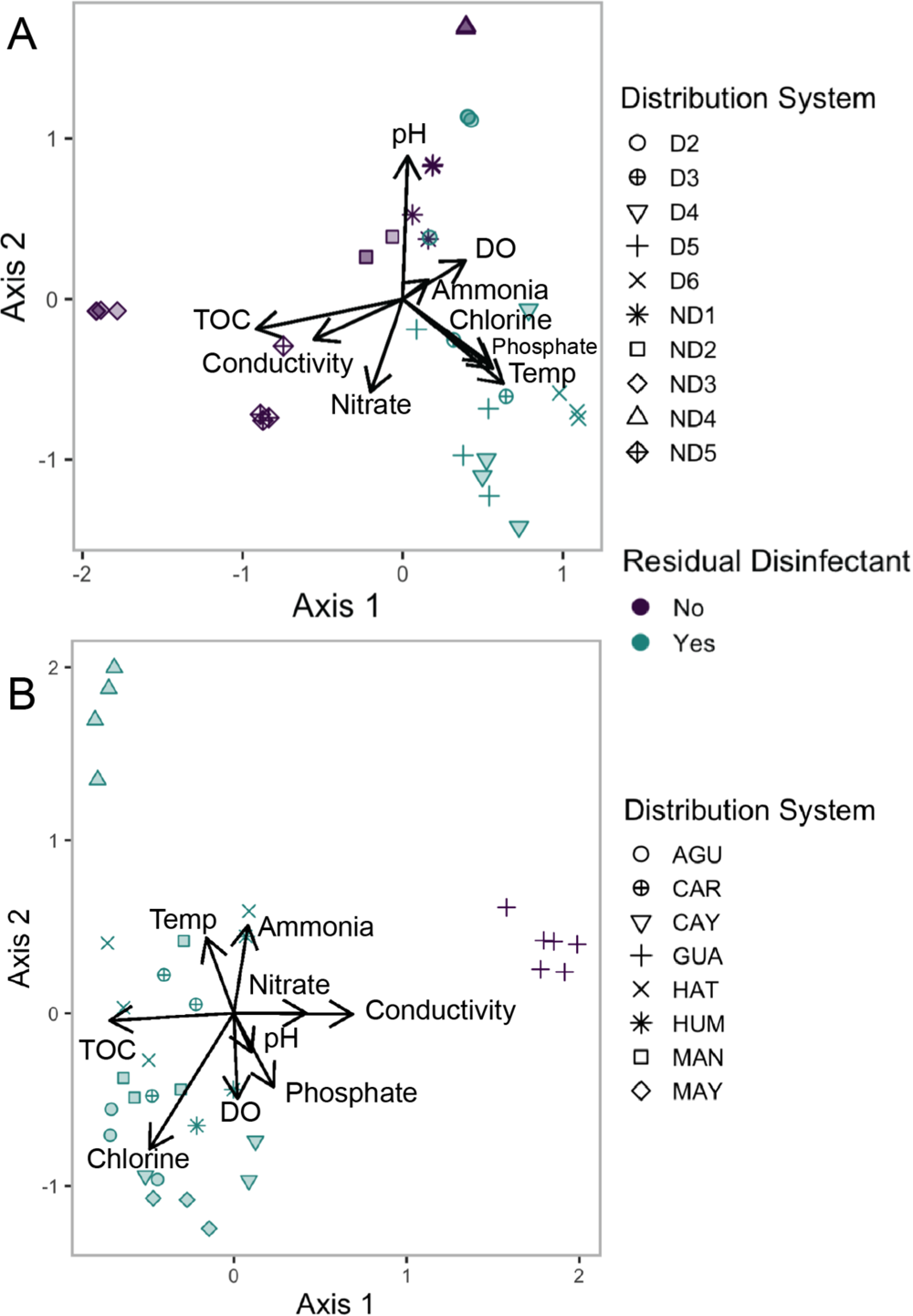
Diversity of viral populations cluster based on residual disinfectant use and water quality. dbRDA ordination of the Bray Curtis dissimilarities of the viral community structure for (A) Dai et al. (2020) and (B) Sevillano et al (2021) samples. Distribution system is represented by shape and residual disinfectant use by color. The ordination is overlaid with the gradient of the fit between the measured environmental parameters and the viral community dissimilarities. Only samples from distribution systems using chlorine in the Cfb Köppen climate zone from the Dai et al. (2020) study were included in 4A.

#### 3.1.3 Systems with residual disinfectant have less diverse and less even viral communities

To assess the effect of residual disinfectant on the viral communities, we first assessed within-sample diversity (alpha diversity). The average normalized viral richness was 350 ± 820 and the average Shannon evenness index was 0.28 ± 0.10. The Shannon dissimilarities for the drinking water viral populations range from 0.26-7.7, which is consistent with ranges reported for other freshwater sources (Aguirre De Cárcer et al., 2015; Tseng et al., 2013). Distribution systems that used a residual disinfectant had less diverse viral communities than distribution systems that did not use a residual disinfectant. This trend held for systems using both chlorine and chloramine, as well as for the suspected residual disinfectant samples (Tukey’s test p_adj_ ≤ 10^-6^; Figure S5).

Further, no difference was found between the chlorine and chloramine samples (Tukey’s test p_adj_ = 0.92), nor between the suspected residual disinfectant samples and either the chlorine (Tukey’s test p_ad j_= 0.25) or chloramine (Tukey’s test p_adj_ = 0.85) samples. Together, these trends supported our prediction that samples from studies with suspected residual disinfectant usage were indeed likely obtained from systems operated with a residual disinfectant.

Bacterial communities in drinking water systems with residual disinfectant have been observed to have lower richness and evenness (Bautista-de los Santos et al., 2016; Dai et al., 2020). Dai et al. (2020) proposed that disinfectant residuals exert a stronger selective pressure than the low phosphate and ammonia availability in distribution systems that do not use a residual disinfectant. For the Dai et al. (2020) samples, we similarly found that viral communities in drinking water distribution systems with a residual disinfectant (i.e., free chlorine) were less diverse and less even than distribution systems without a residual disinfectant (Wilcoxon rank sum test p-values: normalized richness p-value = 3.0*10^-6^ and Shannon evenness index p-value = 0.001; Figure 4). Correlations between bacterial and viral diversity have been observed in a range of environments (Anderson et al., 2017; Dalcin Martins et al., 2018; Yang et al., 2019).

**Figure 4.**
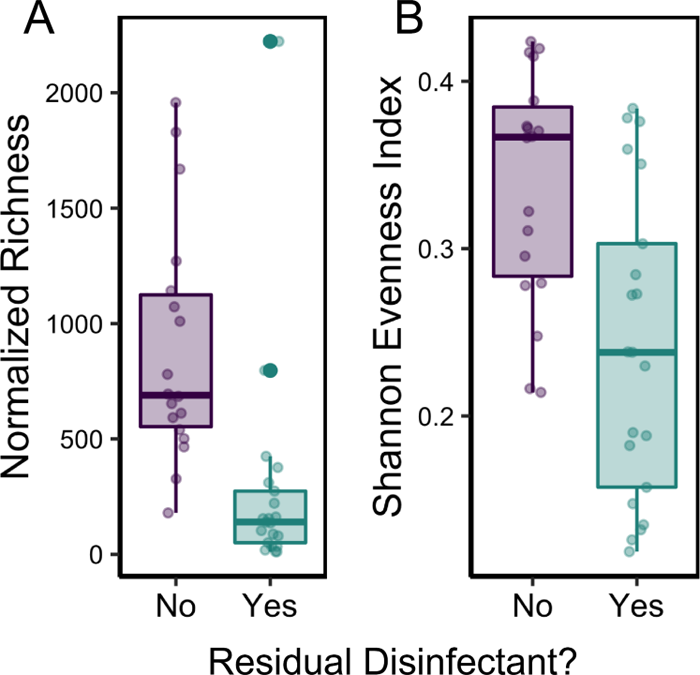
Viral diversity varied based on whether samples originated from distribution systems that use or do not use a residual disinfectant. (A) Observed normalized viral richness and (B) Shannon Evenness Index of the viral community of each sample. Note that only samples from the Cfb Köppen climate zone and samples with chlorine as the residual disinfectant from the Dai et al. (2020) study were included.

Whether residual disinfectant selects for certain viral populations in drinking water, thereby reducing viral community diversity, or the viral population dynamics are following those of their bacterial hosts as they are affected by residual disinfectant and other changing environmental parameters, is unknown. As phages impose a selective pressure on bacterial communities, phages can both disrupt desirable bacterial communities during industrial processes (e.g., yogurt production), as well as be used to prevent contamination by pathogens (e.g., to prevent milk spoilage) (Marks and Sharp, 2000). Based on the abundance and diversity of phages in drinking water, we hypothesize that phages are an important factor influencing the stability of bacterial communities in drinking water. However, more systematic and deeply sequenced analyses of paired cellular and viral metagenomes is needed to better resolve virus, host, and infection dynamics in drinking water distribution systems and their consequences for water quality.

Residual disinfectant is also a significant determinant of between-sample bacterial community structure (Bautista-de los Santos et al., 2016; Dai et al., 2020), e.g., beta diversity. Therefore, we anticipated it to have a similar effect on viral community structure. The viral communities from the Dai et al. (2020) study clustered based on residual disinfectant use (PERMANOVA R^2^=0.09, p-value=0.0001; Figure 3A). While the Dai et al. (2020) samples considered were all taken from the same Köppen climatic zone, the residual disinfectant samples were taken in the summer/fall and have lower conductivities and the no residual disinfectant samples were taken during the winter/spring and have higher conductivities (Tables S2, S10). For this reason, the differences in viral community structure may reflect the additive effect of water quality differences that reflect treatment (e.g., chlorine, phosphate, DO, ammonia) and other factors that may confound the effect of treatment, such as season and water source (e.g., temperature and conductivity). Both season (Pinto et al., 2014; Potgieter et al., 2018; Prest et al., 2016) and water source (Assche et al., 2019; Roeselers et al., 2015) have been shown to influence drinking water microbial communities. When season and water source have been examined together, systems that are groundwater fed have lacked the strong seasonal trends observed in surface water fed systems (Assche et al., 2019; Roeselers et al., 2015). Therefore, season may have a weaker confounding effect than source water for the no residual disinfectant samples, as their conductivities suggest groundwater sources (Douterelo et al., 2018).

Providing an additional opportunity to consider the effect of residual disinfectant, one site from the Sevillano et al. (2021) study, GUA, had chlorine concentrations below the detection limit or much lower than the other sites (chlorine concentration < 0.05 mg/L). The viral community structure of the GUA samples was distinct compared to the other samples (PERMANOVA R^2^=0.18, p-value=0.0002; Figure 3B), including those that had similar conductivities, temperatures, and other water quality parameters. While this further supports the influence of residual disinfectant on viral community structure, more studies are necessary to determine the distinct effects of residual disinfectant use on the viral communities of drinking water distribution systems separate from season and water source.

### 3.2 The metabolic potential of drinking water viromes

Viral genomes often include genes necessary for replication, as well as host-derived auxiliary metabolic genes (AMGs) thought to reprogram host cellular metabolism to support infection processes (Warwick-Dugdale et al., 2019). Characterizing and understanding the influences on the metabolic potential of drinking water viruses is, therefore, an essential step in understanding their influence on the cellular members of the drinking water microbiome.

#### 3.2.1 Water quality parameters are correlated with differences in viral community metabolic potential

The viral community metabolic potential varies with water quality parameters in similar ways to the viral community structure. Specifically, samples clustered primarily by disinfectant status (Figure 5A) and all tested water quality parameters correlated with the metabolic profiles of samples from systems using residual disinfectant, other than DO (Figure 7; PERMANOVA Table S11). Furthermore, the strong correlation between metabolic profiles and drinking water distribution system (PERMANOVA R^2^=0.30, p-value=0.001) and large degree of unexplained variability (residual PERMANOVA R^2^=0.21) suggested that much of the metabolic variability was due to unreported water quality and treatment specific differences. Unlike the viral populations, which were more diverse in distribution systems with no residual disinfectant, there was more diversity in metabolic potential in the residual disinfectant systems (Wilcoxon rank sum test p-value = 0.03; Figure S6). This may reflect an evolutionary benefit of carrying extra genes to survive the oxidative stress of residual disinfectant presence.

**Figure 5:**
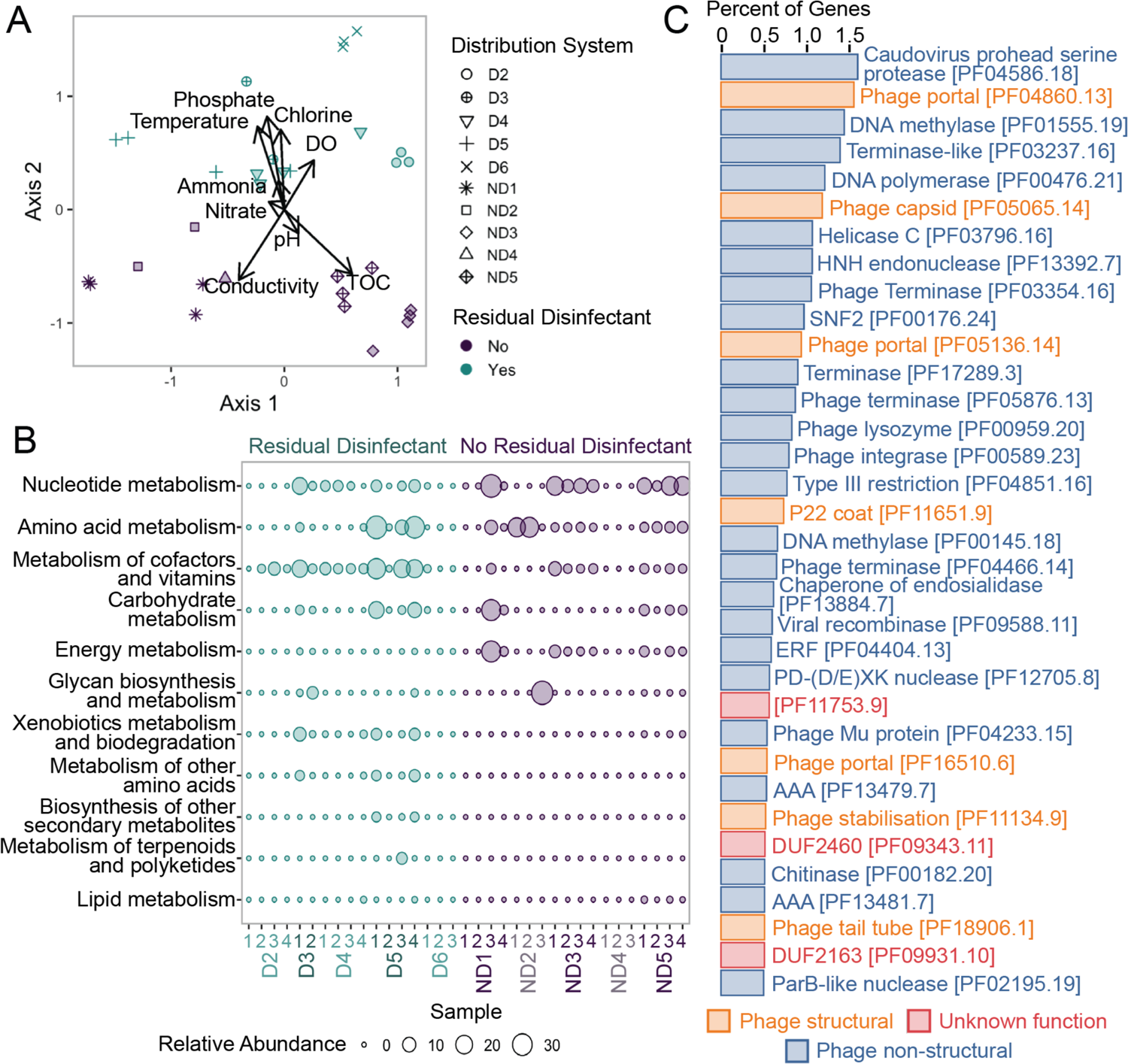
Metabolic potential of the drinking water virome. (A) dbRDA ordination reveals that the diversity of metabolic potential clusters based on residual disinfectant use and water quality. The shape of the points represents the distribution system, and the color represents the residual disinfectant. The ordination is overlaid with the gradient of the fit between the measured environmental parameters and the Bray Curtis dissimilarities. (B) KEGG Ontology (KO) Pathway module relative abundance for the samples from Dai et al (2020). The size of the bubble represents the relative abundance of that pathway module among the reads in that sample. Only samples from the Cfb Köppen climate zone from the Dai et al. (2020) study were included in A and B. (C) Most abundant protein families (Pfam). The percent of genes is the percentage of the number of genes with a specific Pfam annotation divided by the total number of genes with a Pfam annotation. Only those Pfams with a percentage greater than 0.5% are shown. Bars are colored based on whether they were phage non-structural, phage structural, or genes of unknown function. *Figures 6A and 6B* only includes the samples from distribution systems using chlorine and from the Cfb Köppen climate zone.

The relative abundance of KO metabolic pathway modules also differed between distribution systems that used a residual disinfectant and those that did not (Figure 5B, Table S12). Viral contigs from distribution systems that did not use a residual disinfectant were enriched in genes with nucleotide metabolism annotations. Contrastingly, viral contigs from distribution systems that used a residual disinfectant were enriched in genes involving the metabolism of cofactors and vitamins and for xenobiotics biodegradation and metabolism. The higher availability of genes related to these two pathway modules in residual disinfectant systems may reflect the evolutionary pressure to carry genes capable of mitigating the effects of reactive chlorine species (Huang et al., 2021; Kavagutti et al., 2019). However, these trends, particularly the overabundance of nucleotide metabolism in no residual disinfectant systems, could potentially be an artifact of the increased diversity of these samples (Figure 4). As more than 85% of the KOs were found in less than a quarter of the samples (Figure S7), further studies with deeper sequencing or viral-enriched genomes are necessary to robustly investigate differential abundances of molecular functions.

#### 3.2.2 Drinking water viruses encode metabolic genes with the potential to augment microbial metabolic processes

As is common in other environmental viromes (Deboutte et al., 2020; Gregory et al., 2019), most of the predicted viral genes identified in this study were novel, meaning they do not share homology with known genes in public databases. As a result, only 25% had a Pfam annotation and 6% could be assigned a KEGG orthology (KO) molecular function. Of the six most prevalent Pfams, two were phage structural protein families and four were phage non-structural proteins (Figure 5C). The most abundant was a Caudovirus prohead serine protease (PF04586.18), an enzyme involved in capsid maturation. The ubiquity of viral-related annotations among the most abundant Pfams supports that viral contigs were successfully sorted from the majority cellular metagenomes.

The top four most abundant KO metabolic modules were nucleotide metabolism (44% of the reads that mapped to viral genes with KO annotations), cofactors and vitamins metabolism (18%), amino acid metabolism (11%), and carbohydrate metabolism (10%). Genes related to nucleotide and amino acid metabolism are commonly abundant in viral genomes, due to their importance in viral replication (Hurwitz et al., 2015). Carbohydrate metabolism genes carried by viruses may be involved in reprogramming central carbon metabolism to favor energy production necessary for viral replication (Howard-Varona et al., 2020). In the oligotrophic environment of the drinking water microbiome, these genes, along with those encoding for proteins related to cofactor and vitamin metabolism, might contribute to the survival of the bacteriophages’ hosts or enhance viral replication.

Nitrogen metabolism genes are common among drinking water bacteria (Potgieter et al., 2020) to survive the oligotrophic conditions of drinking water (Kooij and Wielen, 2013). In other nitrogen-limited environments, phages have carried more genes related to nitrogen cycling (Warwick-Dugdale et al., 2019). Similarly, genes related to nitrogen metabolism were identified in the drinking water virome in this study (Figure S8), including carbonic anhydrase (K01673), glutamine synthetase (K01915), nitrate reductase/nitrite oxidoreductase (K00370), glutamate dehydrogenase (K15371), nitric oxide reductase (K04748), nitrate/nitrite transporter (K02575), and nitrogen fixation protein NifU (K04488). The most ubiquitous nitrogen metabolism related genes were glutamine synthetase (K01915; 11 assemblies), followed by NifU (K04488; 8 assemblies). In addition to being part of nitrogen metabolism, glutamine synthetase is also involved in amino acid metabolism. The presence of viral encoded nitrogen-related genes suggests that in drinking water distribution systems, viruses may impact nutrient metabolism, either through rewiring host metabolism to support infection or by mediating horizontal gene transfer of nutrient-stress related genes (Warwick-Dugdale et al., 2019; Zimmerman et al., 2020).

To survive in drinking water systems with a residual disinfectant, microorganisms must also mitigate the effects of reactive chlorine species (Gray et al., 2013). In this study, glutathione reductase (K00383) and genes responsible for four of the five steps of coenzyme A biosynthesis (M00120; Figure S9) were identified in the viruses of distribution systems that use a residual disinfectant (59 samples in 5 studies). Low molecular weight thiols, including glutathione and coenzyme A, which maintain the reducing potential of the cytosol (Chesney and Eaton, 1996), have previously been found in bacterial genomes from drinking water distribution systems treated with chlorine (Dai et al., 2020; Douterelo et al., 2018; Gomez-Alvarez et al., 2012). In other environments, genes broadly involved in surviving oxidative stress have been found in phage genomes (Huang et al., 2021; Kavagutti et al., 2019) and have been shown to be beneficial to viral fitness (Holmgren, 1989; Russel and Model, 1986). While viral genes are implicated in combating oxidative stress in distribution systems using a residual disinfectant, additional research is needed to assess the role of viruses in mitigating the harmful effects of reactive chlorine species in drinking water, specifically.

Taken together, the presence of genes related to nitrogen metabolism and oxidative stress mitigation on drinking water viral contigs suggest that phages may contribute to the transfer of genes that are important for microbial survival in drinking water distribution systems.

### 3.3 Opportunities for Future Work

Metagenomic analyses allow researchers to make broad assessments of microbial communities and suggest areas for more targeted studies. The analyses conducted here suggest that residual disinfectant use impacts the diversity and metabolic potential of the drinking water virome.

However, future bench-scale experiments will be necessary to separate the effects of residual disinfectant from other water quality and sampling differences. Confounding methodological differences between studies, such as DNA extraction method, can mask or be correlated with the factors, including climate, geography, and treatment regime, that may shape the microbial communities of drinking water. While standardizing methodologies may be impossible and premature, coordination between researchers could facilitate cross-study comparisons (Bautista-de los Santos et al., 2016; Hull et al., 2019). An excellent first step would be to make raw sequencing data public and report temperature, water quality characteristics (e.g., chlorine, phosphate, ammonia, nitrate, TOC concentrations), and biomass proxies (e.g., ATP levels, cell counts), as well as other metadata (e.g., source water characteristics, details of the treatment, season, location) (Bautista-de los Santos et al., 2016; Yilmaz et al., 2011). Application of standardized spike-ins, such as “sequins” (Hardwick et al., 2018), to drinking water microbiomes would further facilitate between study comparisons, as well as reveal whether the changes in viral gene relative abundances described in this work mirror the trends in absolute abundance.

For the most robust assessments of the drinking water virome, future studies must specifically develop sample processing methods that enrich for viruses and are optimized for the sample matrix (Langenfeld et al., 2021). As the drinking water samples of this meta-analysis were not collected with the intention of enriching for submicron viral particles, the sequenced viruses may be biased towards particle-associated viruses, viruses actively infecting a cellular host, and free viruses trapped by the chosen filter. Future studies designed specifically to query the viral fraction of drinking water (including both DNA and RNA viruses) and with deeper sequencing efforts are necessary to determine whether the trends described here are reflective of the full viral community. To more fully capture both the drinking water viral community and their influences on bacterial communities, future studies should also explore viruses in drinking water biofilms.

As part of this effort, future work should characterize the viral genes that are on DNA within viral particles, as compared to extracellular DNA. Altogether, these studies will also determine the extent to which phage-facilitated metabolic reprogramming or gene transfer contributes to bacterial survival in drinking water systems. By connecting the drinking water virome’s dynamics with that of the bacterial community, improved treatment processes may become possible.

## 4 Conclusions

- Mining drinking water metagenomes for viral sequences reveals differences in the viral community based on geography and water quality parameters.
- More diverse and even viral communities are found in distribution systems that do not use a residual disinfectant.
- Viruses in drinking water carry genes, such as those for nitrogen metabolism and chlorine stress mitigation, that mirror those identified as critical for the drinking water bacterial community.

## Acknowledgements

We would like to thank the Remaking Water Infrastructure Blue Sky team, particularly Sarah Potgieter, Christopher Anderson, and Katherine Dowdell, as well as the Duhaime and Wigginton labs for contributing their technical and intellectual expertise for bioinformatic experimental design and analyses and manuscript preparation. This work was supported by the Blue Sky Initiative of the University of Michigan College of Engineering and NSF CAREER funding to Ameet Pinto [NSF CBET 1749530].

## Notes

### Competing Interest Statement

The authors have declared no competing interest.

## Citations

1. Aguirre De Cárcer, D., López-Bueno, A., Pearce, D.A., Alcamí, A., 2015. Biodiversity and distribution of polar freshwater DNA viruses. Sci. Adv. https://doi.org/10.1126/sciadv.1400127

2. Albinana-Gimenez, N., Clemente-Casares, P., Bofill-Mas, S., Hundesa, A., Ribas, F., Girones, R., 2006. Distribution of Human Polyomaviruses, Adenoviruses, and Hepatitis E Virus in the Environment and in a Drinking-Water Treatment Plant. Environ. Sci. Technol. 40, 7416–7422. https://doi.org/10.1021/es060343i

3. Anderson, C.L., Sullivan, M.B., Fernando, S.C., 2017. Dietary energy drives the dynamic response of bovine rumen viral communities. Microbiome 5, 155. https://doi.org/10.1186/s40168-017-0374-3

4. Van Assche, A., Lievens, B., Crauwels, S., De Brabanter, J., Willems, K.A., 2019. Characterization of the bacterial community composition in water of drinking water production and distribution systems in 1–11. https://doi.org/10.1002/mbo3.726

5. Bautista-de los Santos, Q.M., Schroeder, J.L., Sevillano-Rivera, M.C., Sungthong, R., Ijaz, U.Z., Sloan, W. T, Pinto, A.J., 2016. Emerging investigators series: microbial communities in full-scale drinking water distribution systems – a meta-analysis. Environ. Sci. Water Res 2, 631–644. https://doi.org/10.1039/c6ew00030d

6. Bin Jang, H., Bolduc, B., Zablocki, O., Kuhn, J.H., Roux, S., Adriaenssens, E.M., Brister, J.R., Kropinski, A.M., Krupovic, M., Lavigne, R., Turner, D., Sullivan, M.B., 2019. Taxonomic assignment of uncultivated prokaryotic virus genomes is enabled by gene-sharing networks. Nat. Biotechnol. 37, 632–639. https://doi.org/10.1038/s41587-019-0100-8

7. Bohus, V., Toth, E.M., Szekely, A.J., Makk, J., Baranyi, K., Patek, G., Schunk, J., Marialigeti, K., 2010. Microbiological investigation of an industrial ultra pure supply water plant using cultivation-based and cultivation-independent methods. Water Res. 44, 6124–6132. https://doi.org/10.1016/j.watres.2010.07.006

8. Brinkman, N.E., Villegas, E.N., Garland, J.L., Keely, S.P., 2018. Reducing inherent biases introduced during DNA viral metagenome analyses of municipal wastewater. PLoS One 13, 1–23. https://doi.org/10.1371/journal.pone.0195350

9. Brumfield, K.D., Hasan, N.A., Id, M.B.L., Cotruvo, J.A., Rashed, M., Id, R.R.C., Huq, A., 2020. A comparative analysis of drinking water employing metagenomics. PLoS One 15, 1–27. https://doi.org/10.1371/journal.pone.0231210

10. Chao, Y., Ma, L., Yang, Y., Ju, F., Zhang, X.X., Wu, W.M., Zhang, T., 2013. Metagenomic analysis reveals significant changes of microbial compositions and protective functions during drinking water treatment. Sci. Rep. 3, 1–9. https://doi.org/10.1038/srep03550

11. Chen, S., Zhou, Y., Chen, Y., Gu, J., 2018. fastp: an ultra-fast all-in-one FASTQ preprocessor 884–890. https://doi.org/10.1093/bioinformatics/bty560

12. Chesney, J.A., Eaton, J.W., 1996. Bacterial Glutathione: a Sacrificial Defense against Chlorine Compounds. J. Bacteriol. 178, 2131–2135.

13. Dai, Z., Sevillano-rivera, M.C., Calus, S.T., Santos, Q.M.B.L., Eren, A.M., Van Der Wielen, P.W.J.J., Ijaz, U.Z., Pinto, A.J., 2020. Disinfection exhibits systematic impacts on the drinking water microbiome. Microbiome 1–19. https://doi.org/10.1186/s40168-020-00813-0

14. Dalcin Martins, P., Danczak, R.E., Roux, S., Frank, J., Borton, M.A., Wolfe, R.A., Burris, M.N., Wilkins, M.J., 2018. Viral and metabolic controls on high rates of microbial sulfur and carbon cycling in wetland ecosystems. Microbiome 6, 1–17. https://doi.org/10.1186/s40168-018-0522-4

15. Deboutte, W., Beller, L., Kwe, C., Maes, P., De Graaf, D.C., 2020. Honey-bee – associated prokaryotic viral communities reveal wide viral diversity and a profound metabolic coding potential 117, 10511–10519. https://doi.org/10.1073/pnas.1921859117

16. Delong, E.F., Preston, C.M., Mincer, T., Rich, V., Hallam, S.J., Frigaard, N., Martinez, A., Sullivan, M.B., Edwards, R., Brito, B.R., Chisholm, S.W., Karl, D.M., 2006. Community Genomics Among Stratified Microbial Assemblages in the Ocean’s Interior. Science (80-.). 311, 496–504. https://doi.org/10.1126/science.1120250

17. Dias, R.S., Abe, A.E., Lima, H.S., Silva, L.C.F., de Paula, S.O., da Silva, C.C., 2020. Viral concentration methods for diversity studies in soil samples. Appl. Soil Ecol. 155, 103666. https://doi.org/10.1016/j.apsoil.2020.103666

18. Dong, Y., Kim, J., Lewis, G.D., 2010. Evaluation of methodology for detection of human adenoviruses in wastewater, drinking water, stream water and recreational waters. J. Appl. Microbiol. 108, 800–809. https://doi.org/10.1111/j.1365-2672.2009.04477.x

19. Douterelo, I., Calero-Preciado, C., Soria-Carrasco, V., Boxall, J.B., 2018. Whole metagenome sequencing of chlorinated drinking water distribution systems. Environ. Sci. Water Res. Technol. 4, 2080–2091. https://doi.org/10.1039/c8ew00395e

20. Feazel, L.M., Baumgartner, L.K., Peterson, K.L., Frank, D.N., Harris, J.K., Pace, N.R., 2009. Opportunistic pathogens enriched in showerhead biofilms. Proc. Natl. Acad. Sci. 106, 16393–16399. https://doi.org/10.1073/pnas.0908446106

21. Fuhrman, J.A., Noble, R.T., 1995. Viruses and protists cause similar bacterial mortality in coastal seawater 40, 1236–1242.

22. Gall, A.M., Mari, B.J., Lu, Y., Shisler, J.L., 2015. Waterborne Viruses: A Barrier to Safe Drinking Water. PLOS Pathog. 1–7. https://doi.org/10.1371/journal.ppat.1004867

23. Garner, E., Chen, C., Xia, K., Bowers, J., Engelthaler, D.M., Mclain, J., Edwards, M.A., Pruden, A., 2018. Metagenomic Characterization of Antibiotic Resistance Genes in Full-Scale Reclaimed Water Distribution Systems and Corresponding Potable Systems. Environ. Sci. Technol. 52, 6113–6125. https://doi.org/10.1021/acs.est.7b05419

24. Gomez-Alvarez, V., Revetta, R.P., Domingo, J.W.S., 2012. Metagenomic Analyses of Drinking Water Receiving Different Disinfection Treatments. AEM 78, 6095–6102. https://doi.org/10.1128/AEM.01018-12

25. Gray, M.J., Wholey, W.-Y., Jakob, U., 2013. Bacterial Responses to Reactive Chlorine Species. Annu. Rev. Microbiol. 141–160. https://doi.org/10.1146/annurev-micro-102912-142520.Bacterial

26. Gregory, A.C., Zayed, A.A., Conceição-Neto, N., Temperton, B., Bolduc, B., Alberti, A., Ardyna, M., Arkhipova, K., Carmichael, M., Cruaud, C., Dimier, C., Domínguez-Huerta, G., Ferland, J., Kandels, S., Liu, Y., Marec, C., Pesant, S., Picheral, M., Pisarev, S., Poulain, J., Tremblay, J.É., Vik, D., Acinas, S.G., Babin, M., Bork, P., Boss, E., Bowler, C., Cochrane, G., de Vargas, C., Follows, M., Gorsky, G., Grimsley, N., Guidi, L., Hingamp, P., Iudicone, D., Jaillon, O., Kandels-Lewis, S., Karp-Boss, L., Karsenti, E., Not, F., Ogata, H., Poulton, N., Raes, J., Sardet, C., Speich, S., Stemmann, L., Sullivan, M.B., Sunagawa, S., Wincker, P., Culley, A.I., Dutilh, B.E., Roux, S., 2019. Marine DNA Viral Macro- and Microdiversity from Pole to Pole. Cell 177, 1109–1123.e14. https://doi.org/10.1016/j.cell.2019.03.040

27. Guidi, L., Chaffron, S., Bittner, L., Eveillard, D., Larhlimi, A., Roux, S., Darzi, Y., Audic, S., Berline, L., Brum, J.R., Coelho, L.P., Cesar, J., Espinoza, I., Malviya, S., Sunagawa, S., Dimier, C., Kandels-lewis, S., Picheral, M., Poulain, J., 2016. Plankton networks driving carbon export in the oligotrophic ocean. Nature. https://doi.org/10.1038/nature16942

28. Gulino, K., Rahman, J., Badri, M., Morton, J., Bonneau, R., Ghedin, E., 2020. Initial Mapping of the New York City Wastewater Virome. mSystems 5, 1–18. https://doi.org/10.1128/msystems.00876-19

29. Guo, J., Bolduc, B., Zayed, A.A., Varsani, A., Dominguez-huerta, G., Delmont, T.O., Pratama, A.A., Gazitúa, M.C., Vik, D., Sullivan, M.B., Roux, S., 2021. VirSorter2: a multi-classifier, expert-guided approach to detect diverse DNA and RNA viruses. Microbiome 9, 9–37. https://doi.org/10.1186/s40168-020-00990-y

30. Gurevich, A., Saveliev, V., Vyahhi, N., Tesler, G., 2013. QUAST: quality assessment tool for genome assemblies. Bioinformatics 29, 1072–1075. https://doi.org/10.1093/bioinformatics/btt086

31. Hardwick, S.A., Chen, W.Y., Wong, T., Kanakamedala, B.S., Deveson, I.W., Ongley, S.E., Santini, N.S., Marcellin, E., Smith, M.A., Nielsen, L.K., Lovelock, C.E., Neilan, B.A., Mercer, T.R., 2018. Synthetic microbe communities provide internal reference standards for metagenome sequencing and analysis. Nat. Commun. 9, 1–10. https://doi.org/10.1038/s41467-018-05555-0

32. Holmgren, A., 1989. Electron Transport to Reductive Enzymes 264, 13963–13966.

33. Howard-Varona, C., Lindback, M.M., Bastien, G.E., Solonenko, N., Zayed, A.A., Jang, H., Andreopoulos, B., Brewer, H.M., del Rio, T.G., Adkins, J.N., Paul, S., Sullivan, M.B., Duhaime, M.B., 2020. Phage-specific metabolic reprogramming of virocells. ISME J. 14, 881–895. https://doi.org/10.1038/s41396-019-0580-z

34. Huang, X., Jiao, N., Zhang, R., 2021. The genomic content and context of auxiliary metabolic genes in roseophages. Environ. Microbiol. 23, 3743–3757. https://doi.org/10.1111/1462-2920.15412

35. Hull, N.M., Ling, F., Pinto, A.J., Albertsen, M., Jang, H.G., Hong, P.Y., Konstantinidis, K.T., LeChevallier, M., Colwell, R.R., Liu, W.T., 2019. Drinking Water Microbiome Project: Is it Time? Trends Microbiol. 27, 670–677. https://doi.org/10.1016/j.tim.2019.03.011

36. Hurwitz, B.L., Brum, J.R., Sullivan, M.B., 2015. Depth-stratified functional and taxonomic niche specialization in the ‘core’ and ‘flexible’ Pacific Ocean Virome 472–484. https://doi.org/10.1038/ismej.2014.143

37. Hurwitz, B.L., Deng, L., Poulos, B.T., Sullivan, M.B., 2013. Evaluation of methods to concentrate and purify ocean virus communities through comparative, replicated metagenomics. Environ. Microbiol. 15, 1428–1440. https://doi.org/10.1111/j.1462-2920.2012.02836.x

38. Jia, S., Shi, P., Hu, Q., Li, B., Zhang, T., Zhang, X., 2015. Bacterial Community Shift Drives Antibiotic Resistance Promotion during Drinking Water Chlorination. Environ. Sci. Technol. 49, 12271–12279. https://doi.org/10.1021/acs.est.5b03521

39. Jia, S., Wu, J., Ye, L., Zhao, F., Li, T., Zhang, X., 2019. Metagenomic assembly provides a deep insight into the antibiotic resistome alteration induced by drinking water chlorination and its correlations with bacterial host changes. J. Hazard. Mater. 379, 120841. https://doi.org/10.1016/j.jhazmat.2019.120841

40. Kallies, R., Hölzer, M., Toscan, R.B., Nunes, U., 2019. Evaluation of Sequencing Library Preparation Protocols for Viral Metagenomic Analysis from Pristine Aquifer Groundwaters. Viruses 11. doi:https://doi.org/10.3390/v11060484

41. Kavagutti, V.S., Andrei, A.-Ş., Mehrshad, M., Salcher, M.M., Ghai, R., 2019. Phage-centric ecological interactions in aquatic ecosystems revealed through ultra-deep metagenomics. Microbiome 7, 1–15. https://doi.org/10.1186/s40168-019-0752-0

42. Kieft, K., Zhou, Z., Anantharaman, K., 2020. VIBRANT: Automated recovery, annotation and curation of microbial viruses, and evaluation of viral community function from genomic sequences. Microbiome 8, 1–23. https://doi.org/10.1186/s40168-020-00867-0

43. van der Kooij, D., van der Wielen, P.W.J.J., 2013. Microbial Growth in Drinking Water Supplies. IWA Publishing.

44. Kothari, A., Roux, S., Zhang, H., Prieto, A., Soneja, D., Chandonia, J., Spencer, S., 2021. Ecogenomics of Groundwater Phages Suggests Niche Differentiation Linked to Specific Environmental Tolerance. mSystems 6, 1–12. https://doi.org/10.1128/mSystems.00537-21

45. Kotlarz, N., Raskin, L., Zimbric, M., Errickson, J., Lipuma, J.J., Caverly, L.J., 2019. Retrospective Analysis of Nontuberculous Mycobacterial Infection and Monochloramine Disinfection of Municipal Drinking Water in Michigan. mSphere 4, 1–8. https://doi.org/10.1128/mSphere.00160-19

46. Krishna, K.C.B., Sathasivan, A., Listowski, A., 2020. Influence of treatment processes and disinfectants on bacterial community compositions and opportunistic pathogens in a full-scale recycled water distribution system. J. Clean. Prod. 274, 123034. https://doi.org/10.1016/j.jclepro.2020.123034

47. Langenfeld, K., Chin, K., Roy, A., Wigginton, K., Duhaime, M.B., 2021. Comparison of ultrafiltration and iron chloride flocculation in the preparation of aquatic viromes from contrasting sample types. PeerJ 9:e11111. https://doi.org/10.7717/peerj.11111

48. Langmead, B., Salzberg, S.L., 2012. Fast gapped-read alignment with Bowtie 2. Nat. Methods 9, 357–9. https://doi.org/10.1038/nmeth.1923

49. Leinonen, R., Sugawara, H., Shumway, M., 2011. The Sequence Read Archive 39, 2010–2012. https://doi.org/10.1093/nar/gkq1019

50. Li, H., Handsaker, B., Wysoker, A., Fennell, T., Ruan, J., Homer, N., Marth, G., Abecasis, G., Durbin, R., Subgroup, 1000 Genome Project Data Processing, 2009. The Sequence Alignment/Map format and SAMtools. Bioinformatics 25, 2078–2079. https://doi.org/10.1093/bioinformatics/btp352

51. Liao, Y., Smyth, G.K., Shi, W., 2014. featureCounts: an efficient general purpose program for assigning sequence reads to genomic features. Bioinformatics 30, 923–930. https://doi.org/10.1093/bioinformatics/btt656

52. Liao, Y., Smyth, G.K., Shi, W., 2013. The Subread aligner: fast, accurate and scalable read mapping by seed-and-vote. Nucleic Acids Res. 41. https://doi.org/10.1093/nar/gkt214

53. Liu, G., Verberk, J.Q.J.C., Van Dijk, J.C., 2013. Bacteriology of drinking water distribution systems: An integral and multidimensional review. Appl. Microbiol. Biotechnol. 97, 9265– 9276. https://doi.org/10.1007/s00253-013-5217-y

54. Liu, L., Xing, X., Hu, C., Wang, H., Lyu, L., 2019. Chemosphere Effect of sequential UV / free chlorine disinfection on opportunistic pathogens and microbial community structure in simulated drinking water distribution systems. Chemosphere 219, 971–980. https://doi.org/10.1016/j.chemosphere.2018.12.067

55. Ma, L., Li, B., Jiang, X.T., Wang, Y.L., Xia, Y., Li, A.D., Zhang, T., 2017. Catalogue of antibiotic resistome and host-tracking in drinking water deciphered by a large scale survey. Microbiome 5, 154. https://doi.org/10.1186/s40168-017-0369-0

56. Ma, L., Li, B., Zhang, T., 2019. New insights into antibiotic resistome in drinking water and management perspectives: A metagenomic based study of small-sized microbes. Water Res. 152, 191–201. https://doi.org/10.1016/j.watres.2018.12.069

57. Malki, K., Rosario, K., Sawaya, N.A., Székely, A.J., Tisza, M.J., Breitbart, M., 2020. Prokaryotic and Viral Community Composition of Freshwater Springs in Florida, USA. mBiom 11, e00436–20. https://doi.org/10.1128/mBio.00436-20

58. Marks, T., Sharp, R., 2000. Bacteriophages and biotechnology: a review. J. Chem. Technol. Biotechnol. 17, 6–17.

59. Mellbye, B.L., Bottomley, P.J., Sayavedra-soto, L.A., 2015. Nitrite-Oxidizing Bacterium Nitrobacter winogradskyi Produces N-Acyl-Homoserine Lactone Autoinducers. AEM. https://doi.org/10.1128/AEM.01103-15

60. Moon, K., Kim, S., Kang, I., Cho, J., 2020. Viral metagenomes of Lake Soyang, the largest freshwater lake in South Korea. Sci. Data 1–6. https://doi.org/10.1038/s41597-020-00695-9

61. Nayfach, S., Camargo, A.P., Schulz, F., Eloe-fadrosh, E., Roux, S., Kyrpides, N.C., 2020. CheckV assesses the quality and completeness of metagenome-assembled viral genomes. Nat. Biotechnol. https://doi.org/10.1038/s41587-020-00774-7

62. NCBI, N.C. for B.I., 2019. BLAST. NCBI, N.C. for B.I., 2016. UniVec.

63. Nurk, S., Meleshko, D., Korobeynikov, A., Pevzner, P.A., 2017. metaSPAdes: a new versatile metagenomic assembler 824–834. https://doi.org/10.1101/gr.213959.116.4

64. Oksanen, J., Blanchet, F.G., Friendly, M., Kindt, R., Legendre, P., McGlinn, D., Wagner, P.R.M., O’Hara, R.B., Simpson, G.L., Solymos, P., Stevens, M.H.H., Szoecs, E., Wagner, H., 2020. vegan: Community Ecology Package.

65. Paez-Espino, D., Eloe-Fadrosh, E.A., Pavlopoulos, G.A., Thomas, A.D., Huntemann, M., Mikhailova, N., Rubin, E., Ivanova, N.N., Kyrpides, N.C., 2016. Uncovering Earth’s virome. Nature 536, 425–430. https://doi.org/10.1038/nature19094

66. Parmar, K.M., Gaikwad, S.L., Dhakephalkar, P.K., Kothari, R., Singh, R.P., 2017. Intriguing interaction of bacteriophage-host association: An understanding in the era of omics. Front. Microbiol. 8. https://doi.org/10.3389/fmicb.2017.00559

67. Percival, S.L., Wyn-Jones, P., 2013. Methods for the Detection of Waterborne Viruses, Second Edi. ed, Microbiology of Waterborne Diseases: Microbiological Aspects and Risks: Second Edition. Elsevier. https://doi.org/10.1016/B978-0-12-415846-7.00022-6

68. Petrovich, M.L., Zilberman, A., Kaplan, A., Eliraz, G.R., Wang, Y., Langenfeld, K., Duhaime, M., Wigginton, K., Poretsky, R., Avisar, D., Wells, G.F., 2020. Microbial and Viral Communities and Their Antibiotic Resistance Genes Throughout a Hospital Wastewater Treatment System. Front. Microbiol. 11, 1–13. https://doi.org/10.3389/fmicb.2020.00153

69. Pinto, A., Schroeder, J., Lunn, M., Sloan, W., Raskin, L., 2014. Spatial-Temporal Survey and Occupancy-Abundance Modeling To Predict Bacterial Community Dynamics in the Drinking Water Microbiome. MBio 5, e01135–14. https://doi.org/10.1128/mBio.01135-14

70. Pinto, A.J., Xi, C., Raskin, L., 2012. Bacterial community structure in the drinking water microbiome is governed by filtration processes. Environ. Sci. Technol. 46, 8851–8859. https://doi.org/10.1021/es302042t

71. Potgieter, S., Pinto, A., Sigudu, M., du Preez, H., Ncube, E., Venter, S., 2018. Long-term spatial and temporal microbial community dynamics in a large-scale drinking water distribution system with multiple disinfectant regimes. Water Res. 139, 406–419. https://doi.org/10.1016/j.watres.2018.03.077

72. Potgieter, S.C., Dai, Z., Venter, S.N., Sigudu, M., Pinto, A.J., 2020. Microbial Nitrogen Metabolism in Chloraminated Drinking. mSphere2 5. https://doi.org/10.1128/mSphere.00274-20. Editor

73. Prest, E.I., Weissbrodt, D.G., Hammes, F., Van Loosdrecht, M.C.M., Vrouwenvelder, J.S., 2016. Long-Term Bacterial Dynamics in a Full-Scale Drinking Water Distribution System. PLoS One 11, 1–20. https://doi.org/10.1371/journal.pone.0164445

74. Rao, C., Waghmare, S., Lakhe, S., 1981. Detection of viruses in drinking water by concentration on magnetic iron oxide. Appl. Environ. Microbiol. 42, 421–426.

75. Ren, J., Song, K., Deng, C., Ahlgren, N.A., Fuhrman, J.A., Li, Y., Xie, X., Sun, F., 2018. Identifying viruses from metagenomic data by deep learning. Quant. Biol. 8, 64–77. https://doi.org/10.1007/s40484-019-0187-4

76. Roeselers, G., Coolen, J., Van Der Wielen, P.W.J.J., Jaspers, M.C., Atsma, A., De Graaf, B., Schuren, F., 2015. Microbial biogeography of drinking water: patterns in phylogenetic diversity across space and time. Environ. Microbiol. 17, 2505–2514. https://doi.org/10.1111/1462-2920.12739

77. Rohwer, F., Prangishvili, D., Lindell, D., 2009. Roles of viruses in the environment. Environ. Microbiol. 11, 2771–2774. https://doi.org/10.1111/j.1462-2920.2009.02101.x

78. Roux, S., Adriaenssens, E.M., Dutilh, B.E., Koonin, E. V, Kropinski, A.M., Krupovic, M., Kuhn, J.H., Lavigne, R., Brister, J.R., Varsani, A., Amid, C., Aziz, R.K., Bordenstein, S.R., Bork, P., Breitbart, M., Cochrane, G.R., Daly, R.A., Desnues, C., Duhaime, M.B., Emerson, J.B., Enault, F., Fuhrman, J.A., Hingamp, P., Hugenholtz, P., Hurwitz, B.L., Ivanova, N.N., Labonté, J.M., Lee, K., Malmstrom, R.R., Martinez-garcia, M., Mizrachi, I.K., Ogata, H., Rodriguez-valera, F., Rosario, K., Schriml, L., Schulz, F., Steward, G.F., Sullivan, M.B., Sunagawa, S., Suttle, C.A., Temperton, B., Tringe, S.G., Thurber, R.V., Webster, N.S., Whiteson, K.L., Wilhelm, S.W., Wommack, K.E., Woyke, T., Wrighton, K.C., Yilmaz, P., Yoshida, T., 2019. Minimum Information about an Uncultivated Virus Genome (MIUViG). Nat. Biotechnol. 37, 29–37. https://doi.org/10.1038/nbt.4306

79. Roux, S., Bolduc, B., 2017. stampede-clustergenomes. https://doi.org/bitbucket.org/MAVERICLab/stampede-clustergenomes/src/master/

80. Roux, S., Brum, J.R., Dutilh, B.E., Sunagawa, S., Duhaime, M.B., Loy, A., Poulos, B.T., Solonenko, N., Lara, E., Poulain, J., Pesant, S., Kandels-lewis, S., Dimier, C., Picheral, M., Searson, S., Cruaud, C., Alberti, A., Duarte, C.M., Gasol, J.M., Vaqué, D., Tara Oceans Coordinators, Bork, P., Acinas, S.G., Wincker, P., Sullivan, M.B., 2016. Ecogenomics and potential biogeochemical impacts of globally abundant ocean viruses. Nat. Publ. Gr. 537, 689–693. https://doi.org/10.1038/nature19366

81. Roux, S., Enault, F., Hurwitz, B.L., Sullivan, M.B., 2015a. VirSorter: Mining viral signal from microbial genomic data. PeerJ 2015, 1–20. https://doi.org/10.7717/peerj.985

82. Roux, S., Enault, F., Robin, A., Ravet, V., Personnic, S., Theil, S., Colombet, J., Sime-Ngando, T., Debroas, D., 2012. Assessing the Diversity and Specificity of Two Freshwater Viral Communities through Metagenomics. PLoS One 7. https://doi.org/10.1371/journal.pone.0033641

83. Roux, S., Hallam, S.J., Woyke, T., Sullivan, M.B., 2015b. Viral dark matter and virus – host interactions resolved from publicly available microbial genomes. Elife 1–20. https://doi.org/10.7554/eLife.08490

84. Russel, M., Model, P., 1986. The Role of Thioredoxin in Filamentous Phage Assembly. J. Biol. Chem. 261, 14997–15005. https://doi.org/10.1016/S0021-9258(18)66819-X

85. Samhan, F.A., Kronlein, M.R., Fakher, U., Kronlein, C., Robert, D., Hashsham, S.A., 2015. Detection and Occurrence of Indicator Organisms and Pathogens 87, 883–900. https://doi.org/10.2175/106143015X14338845155147

86. Saparbaevna, M., Aizhan, A., Turmagambetova, S., Gennadievich, P., Andrey, A., Bogoyavlenskiy, P., Berezin, V.E., 2017. Comparative study of viromes from freshwater samples of the Ile-Balkhash region of Kazakhstan captured through metagenomic analysis. VirusDisease 28, 18–25. https://doi.org/10.1007/s13337-016-0353-5

87. Sevillano, M., Vosloo, S., Cotto, I., Dai, Z., Jiang, T., Santiago, J.M., Padilla, I.Y., Rosario-pabon, Z., Velez, C., Cordero, F., Alshawabkeh, A., Gu, A., Pinto, A.J., 2021. Spatial-temporal targeted and non-targeted surveys to assess microbiological composition of drinking water in Puerto Rico following Hurricane Maria 13. https://doi.org/10.1016/j.wroa.2021.100123

88. Shaffer, M., Borton, M.A., McGivern, B.B., Zayed, A.A., La Rosa, S.L. 0003 3527 8101, Solden, L.M., Liu, P., Narrowe, A.B., Rodríguez-Ramos, J., Bolduc, B., Gazitúa, M.C., Daly, R.A., Smith, G.J., Vik, D.R., Pope, P.B., Sullivan, M.B., Roux, S., Wrighton, K.C., 2020. DRAM for distilling microbial metabolism to automate the curation of microbiome function. Nucleic Acids Res. 48, 8883–8900. https://doi.org/10.1093/nar/gkaa621

89. Shannon, P., Markiel, A., Ozier, O., Baliga, N.S., Wang, J.T., Ramage, D., Amin, N., Schwikowski, B., Ideker, T., 2003. Cytoscape: A Software Environment for Integrated Models of Biomolecular Interaction Networks. Genome Res. 2498–2504. https://doi.org/10.1101/gr.1239303.metabolite

90. Shi, P., Jia, S., Zhang, X., Zhang, T., Cheng, S., Li, A., 2012. Metagenomic insights into chlorination effects on microbial antibiotic resistance in drinking water. Water Res. 47, 111–120. https://doi.org/10.1016/j.watres.2012.09.046

91. Skvortsov, T., De Leeuwe, C., Quinn, J.P., Mcgrath, J.W., Christopher, C., Allen, R., Mcelarney, Y., Watson, C., Arkhipova, K., Lavigne, R., 2016. Metagenomic Characterisation of the Viral Community of Lough Neagh, the Largest Freshwater Lake in Ireland. PLoS One 11, e0140361. https://doi.org/10.1371/journal.pone.0150361

92. Stamps, B.W., Leddy, M.B., Plumlee, M.H., Hasan, N.A., Colwell, R.R., Spear, J.R., 2018. Characterization of the microbiome at the world’s largest potable water reuse facility. Front. Microbiol. 9, 1–16. https://doi.org/10.3389/fmicb.2018.02435

93. Stanish, L.F., Hull, N.M., Robertson, C.E., Harris, J.K., Stevens, J., Spear, J.R., Pace, N.R., 2016. Factors Influencing Bacterial Diversity and Community Composition in Municipal Drinking Waters in the Ohio River Basin, USA. PLoS One 11, 1–21. https://doi.org/10.1371/journal.pone.0157966

94. Suttle, C.A., 2007. Marine viruses — major players in the global ecosystem. Nat. Rev. Microbiol. 5. https://doi.org/10.1038/nrmicro1750

95. Team, R.C., 2021. R: A language and environment for statistical computing. https://doi.org/www.r-project.org/

96. Team, Rs., 2020. RStudio: Integrated Development for R. https://doi.org/www.rstudio.com/

97. Tseng, C.H., Chiang, P.W., Shiah, F.K., Chen, Y.L., Liou, J.R., Hsu, T.C., Maheswararajah, S., Saeed, I., Halgamuge, S., Tang, S.L., 2013. Microbial and viral metagenomes of a subtropical freshwater reservoir subject to climatic disturbances. ISME J. 7, 2374–2386. https://doi.org/10.1038/ismej.2013.118

98. Tsukatani, Y., Hirose, Y., Harada, J., Misawa, N., Mori, K., Inoue, K., 2015. Complete Genome Sequence of the Bacteriochlorophyll b-Producing Photosynthetic Bacterium Blastochloris viridis. Genome Announc. 3, 5–6. https://doi.org/10.1128/genomeA.01006-15.Copyright

99. Vaerewijck, M.J.M., Huys, G., Carlos, J., Swings, J., 2005. Mycobacteria in drinking water distribution systems: ecology and significance for human health. FEMS Microbiol. Rev. 29, 911–934. https://doi.org/10.1016/j.femsre.2005.02.001

100. Wang, H., Proctor, C.R., Edwards, M.A., Pryor, M., Domingo, J.W.S., Ryu, H., Camper, A.K., Olson, A., Pruden, A., 2014. Microbial Community Response to Chlorine Conversion in a Chloraminated Drinking Water Distribution System. ES&T 48, 10624–10633. https://doi.org/10.1021/es502646d

101. Warwick-Dugdale, J., Buchholz, H.H., Allen, M.J., Temperton, B., 2019. Host-hijacking and planktonic piracy: how phages command the microbial high seas. Virol. J. 1, 1–13. https://doi.org/10.1186/s12985-019-1120-1

102. Williamson, S.J., Allen, L.Z., Lorenzi, H.A., Fadrosh, D.W., Brami, D., Thiagarajan, M., McCrow, J.P., Tovchigrechko, A., Yooseph, S., Venter, J.C., 2012. Metagenomic Exploration of Viruses throughout the Indian Ocean. PLoS One 7. https://doi.org/10.1371/journal.pone.0042047

103. Yang, Y., Gu, X., Te, S.H., Goh, S.G., Mani, K., He, Y., Gin, K.Y.H., 2019. Occurrence and distribution of viruses and picoplankton in tropical freshwater bodies determined by flow cytometry. Water Res. 149, 342–350. https://doi.org/10.1016/j.watres.2018.11.022

104. Ye, X.Y., Xing, M., Zhang, Y.L., Xiao, W.Q., Huang, X.N., Cao, Y.G., Gu, K.D., 2012. Real-Time PCR Detection of Enteric Viruses in Source Water and Treated Drinking Water in Wuhan, China. Curr. Microbiol. 65, 244–253. https://doi.org/10.1007/s00284-012-0152-1

105. Yilmaz, P., Kottmann, R., Field, D., Knight, R., Cole, J.R., Amaral-zettler, L., Gilbert, J.A., Karsch-mizrachi, I., Johnston, A., Cochrane, G., Vaughan, R., Hunter, C., Park, J., Morrison, N., Rocca-serra, P., Sterk, P., Arumugam, M., Bailey, M., Baumgartner, L., Birren, B.W., Blaser, M.J., Bonazzi, V., Booth, T., Bork, P., Bushman, F.D., Buttigieg, P.L., Chain, P.S.G., Charlson, E., Costello, E.K., Huot-creasy, H., Dawyndt, P., Desantis, T., Fierer, N., Fuhrman, J.A., Gallery, R.E., Gevers, D., Gibbs, R.A., Gil, I.S., Gonzalez, A., Gordon, J.I., Guralnick, R., Hankeln, W., Highlander, S., Hugenholtz, P., Jansson, J., Kau, A.L., Kelley, S.T., Kennedy, J., Knights, D., Koren, O., Kuczynski, J., Kyrpides, N., Larsen, R., Lauber, C.L., Legg, T., Ley, R.E., Lozupone, C.A., Ludwig, W., Lyons, D., Maguire, E., Methé, B.A., Meyer, F., Muegge, B., Nakielny, S., Nelson, K.E., Nemergut, D., Neufeld, J.D., Newbold, L.K., Oliver, A.E., Pace, N.R., Palanisamy, G., Peplies, J., Ravel, J., Relman, D.A., Assunta-sansone, S., Schloss, P.D., Schriml, L., Sinha, R., Smith, M.I., Sodergren, E., Spor, A., Stombaugh, J., Tiedje, J.M., Ward, D. V, Weinstock, G.M., Wendel, D., White, O., Whiteley, A., Wilke, A., Wortman, J.R., Yatsunenko, T., Glöckner, F.O., 2011. Minimum information about a marker gene sequence (MIMARKS) and minimum information about any (x) sequence (MIxS) specifications. Nat. Biotechnol. 29, 415–420. https://doi.org/10.1038/nbt.1823

106. Zaouri, N., Jumat, M.R., Cheema, T., Hong, P.Y., 2020. Metagenomics-based evaluation of groundwater microbial profiles in response to treated wastewater discharge. Environ. Res. 180, 108835. https://doi.org/10.1016/j.envres.2019.108835

107. Zhang, H., Chang, F., Shi, P., Ye, L., Zhou, Q., Pan, Y., Li, A., 2019. Antibiotic Resistome Alteration by Different Disinfection Strategies in a Full-Scale Drinking Water Treatment Plant Deciphered by Metagenomic Assembly. Environ. Sci. Technol. 53, 2141–2150. https://doi.org/10.1021/acs.est.8b05907

108. Zimmerman, A.E., Varona, C.H.-, Needham, D.M., John, S.G., Worden, A.Z., Sullivan, M.B., Waldbauer, J.R., Coleman, M.L., 2020. Metabolic and biogeochemical consequences of viral infection in aquatic ecosystems. Nat. Rev. Microbiol. 18. https://doi.org/10.1038/s41579-019-0270-x

