## Supplementary material for "A Snapshot of the Global Drinking Water Virome: Diversity and Metabolic Potential Vary with Residual Disinfectant Use": SI Figures

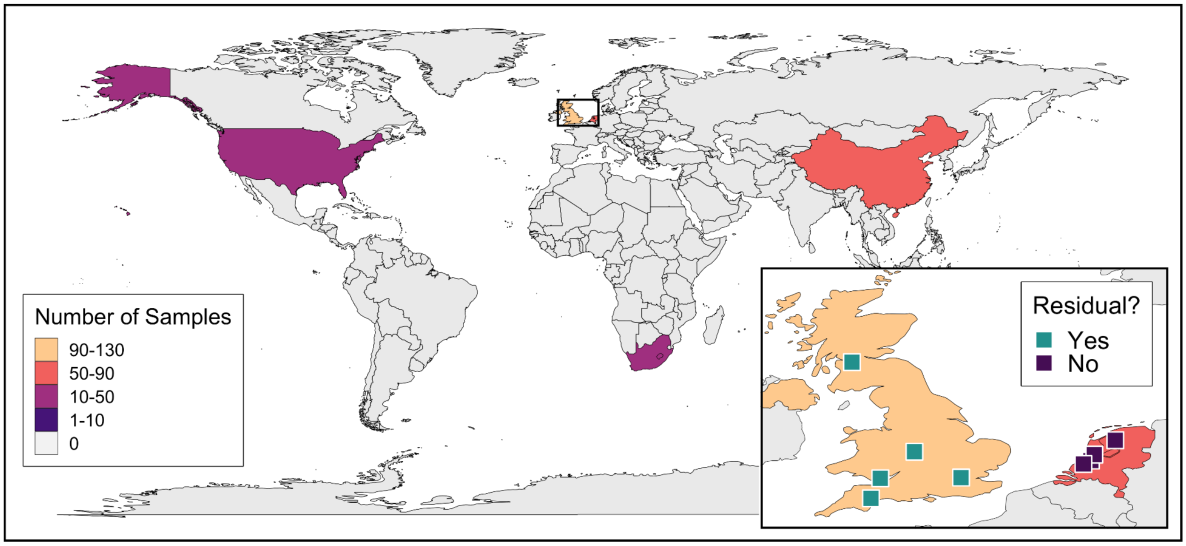


*Figure S1: Depiction of the number of drinking water distribution system samples per country included in this meta-analysis. The color of the country represents the sample number bin. Two samples taken from Singapore are not visible due to the map's scale. Countries with no drinking water metagenomic studies are in grey. Inset map illustrates the locations of the distribution system samples from the Dai et al. (2020) study that were used for subsequent analyses of the differences based on distribution system residual disinfectant use.*

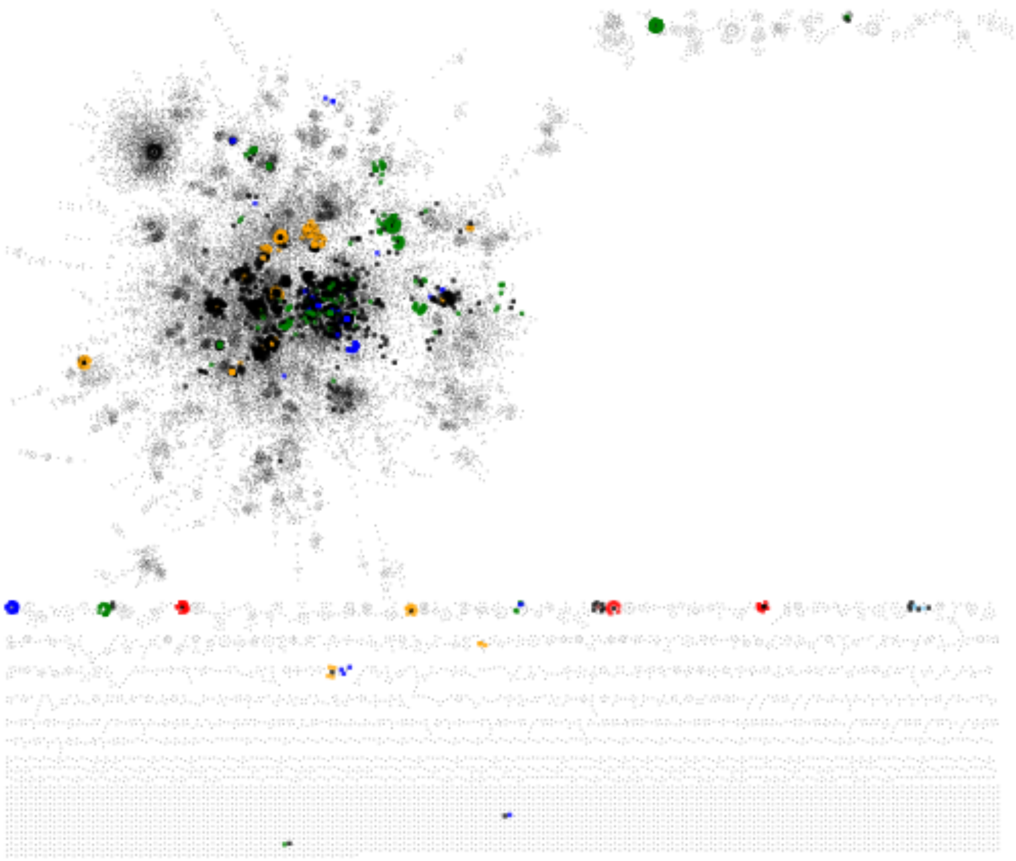

*Figure S2: Network illustration of all viral populations and connected viral genomes with taxonomic annotations from vConTACT2. Viral genomes from the reference database are colored based on their taxonomic assignment (family): Siphoviridae (green), Myoviridae (blue), Podoviridae (orange), Microviridae (red). Black contigs represent drinking water viral populations identified in this study.*

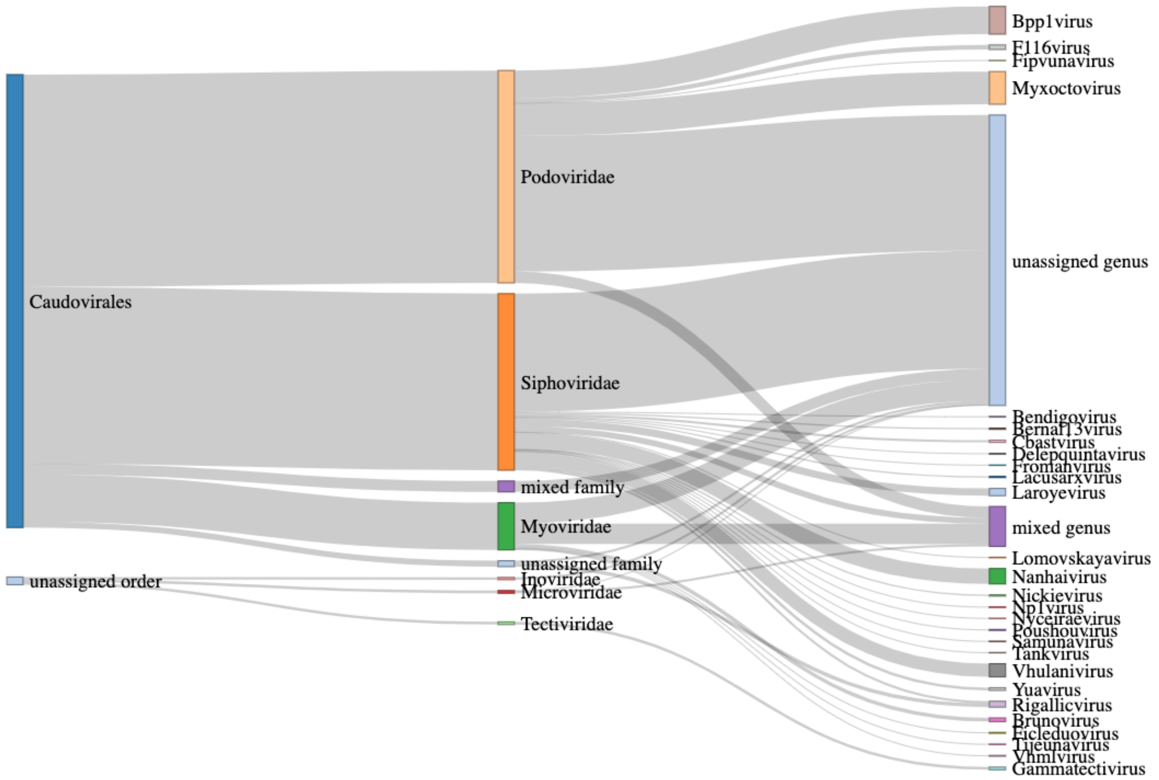


*Figure S3 - Sankey diagram of the taxonomy (from left to right: order, family, genus) of the 320 drinking water viral populations that clustered with known viruses by vConTACT2. 91 viral populations clustered into twenty-five genus level groups, 320 viral populations clustered into six family level groups, and 312 viral populations were assigned to the order Caudovirales. Caudovirales dominates the vConTACT2 database (2,356 out of 2,616 viral genomes) (Bin Jang et al., 2019). Viral populations that were part of a cluster with multiple conflicting reference viruses (family and genus) are designated as "mixed genus" or "mixed family".*

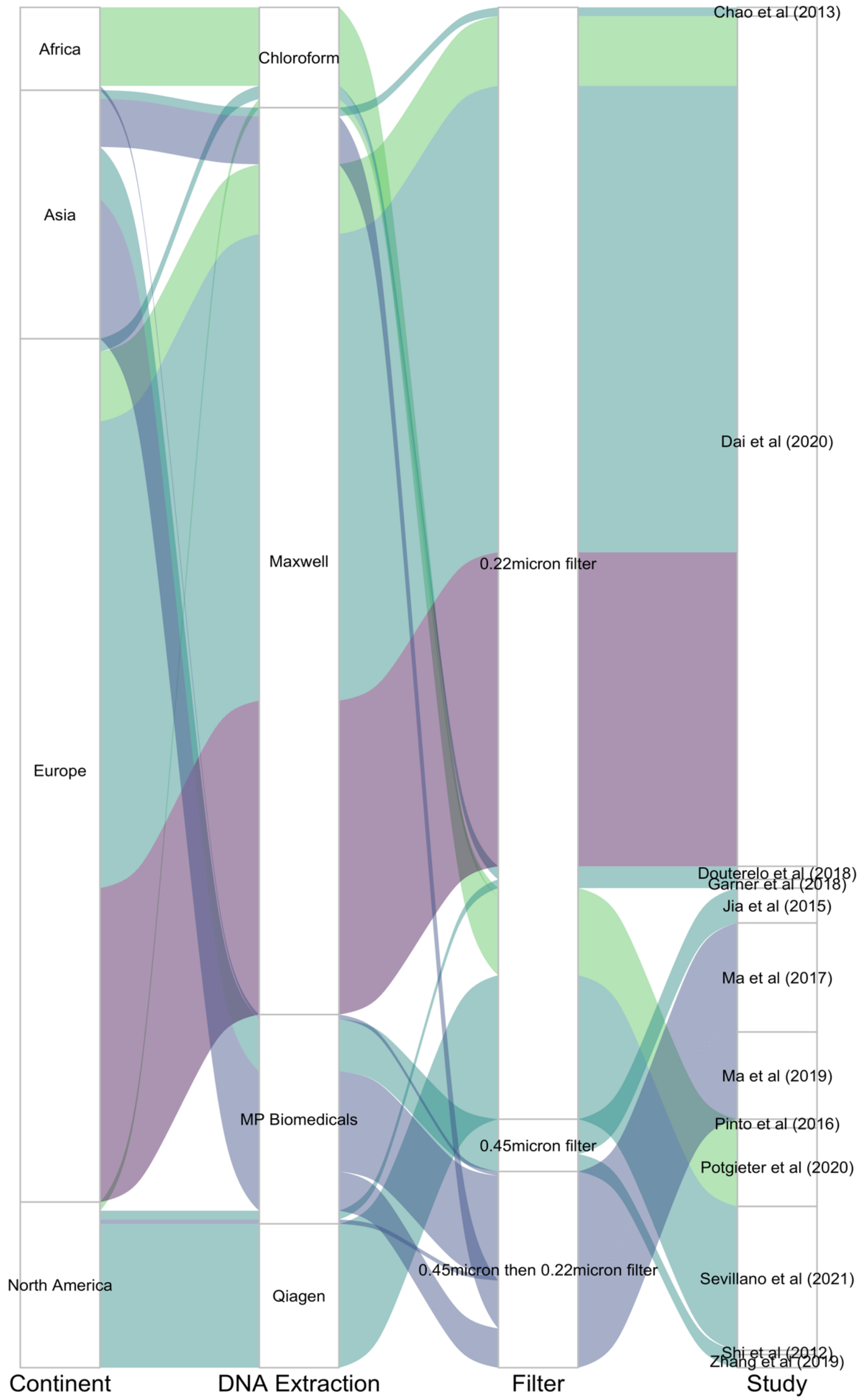

Africa
Asia
Europe
North America
Chloroform
Maxwell
MP Biomedicals
Qiagen
0.22micron filter
0.45micron filter
0.45micron then 0.22micron filter
Chao et al (2013)
Dai et al (2020)
Douterelo et al (2018)
Garner et al (2018)
Jia et al (2015)
Ma et al (2017)
Ma et al (2019)
Pinto et al (2016)
Potgieter et al (2020)
Sevillano et al (2021)
Shi et al (2012)
Zhang et al (2019)
Continent
DNA Extraction
Filter
Study

*Figure S4 - Breakdown of samples based on continent, DNA extraction kit (Chloroform, Maxwell, MP Biomedical, Qiagen), filter used for collecting biomass (0.22 µm, 0.45 µm then 0.22 µm, 0.45 µm), and study (Chao et al., 2013; Dai et al., 2020; Douterelo et al., 2018; Garner et al., 2018; Jia et al., 2015; Ma et al., 2017, 2019; Potgieter et al., 2020; Sevillano et al., 2021; Shi et al., 2012; Zhang et al., 2019). Flows between categories represent the residual disinfectant: samples that used a residual disinfectant (chlorine - teal; chloramines - green; suspected - blue), did not use a residual disinfectant (purple). The metadata for each individual sample can be found in Table S1.*

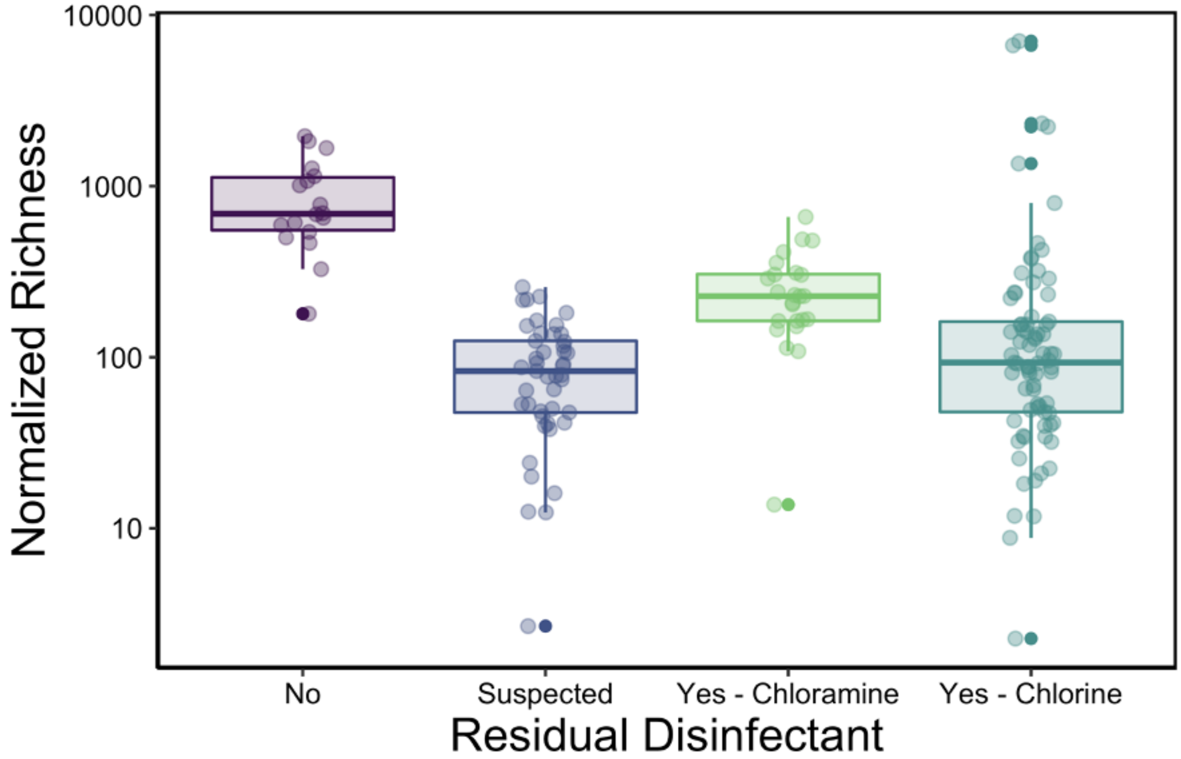


*Figure S5 - Alpha diversity of all samples based on residual disinfectant use. The normalized richness (number of taxa per 100,000 reads) from samples from distribution systems with suspected residual disinfectant, chlorine, and chloramine use are all statistically different from the number of taxa from samples from distribution systems with no residual disinfectant use, while there was no difference in number of taxa between the samples from distribution systems with suspected residual disinfectant, chlorine, and chloramine use.*

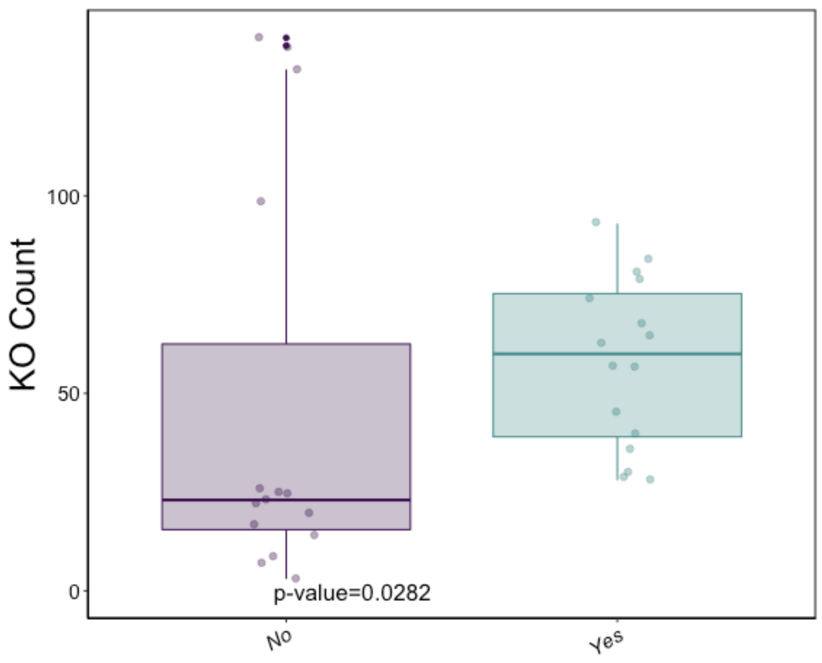


*Figure S6 - Boxplot of the number of KEGGs in a sample overlaid with the individual samples' counts (p-value=0.03).*

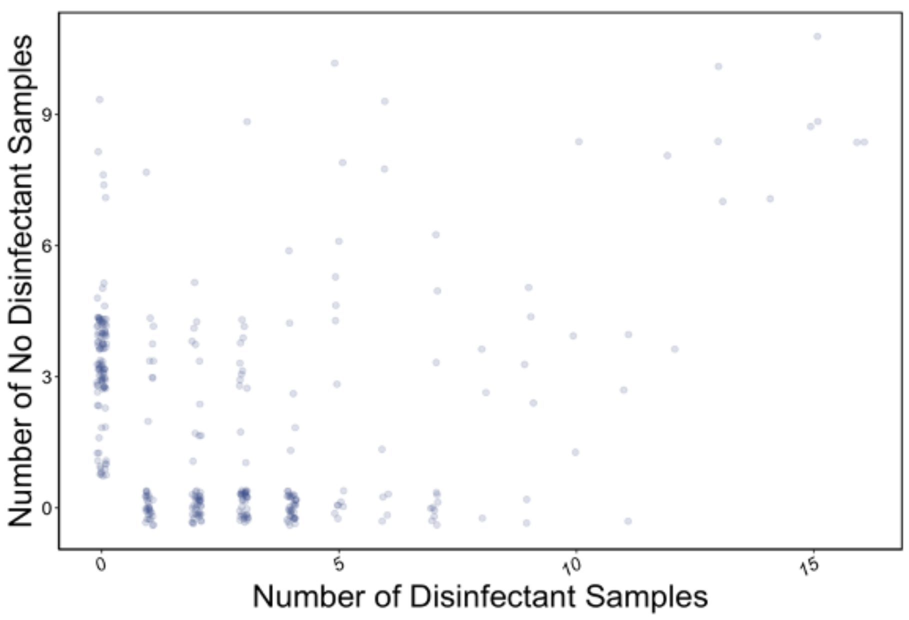


*Figure S7 - Scatterplot of the number of no residual disinfectant samples a given KEGG is in verus the number of residual disinfectant samples that KEGG is in.*

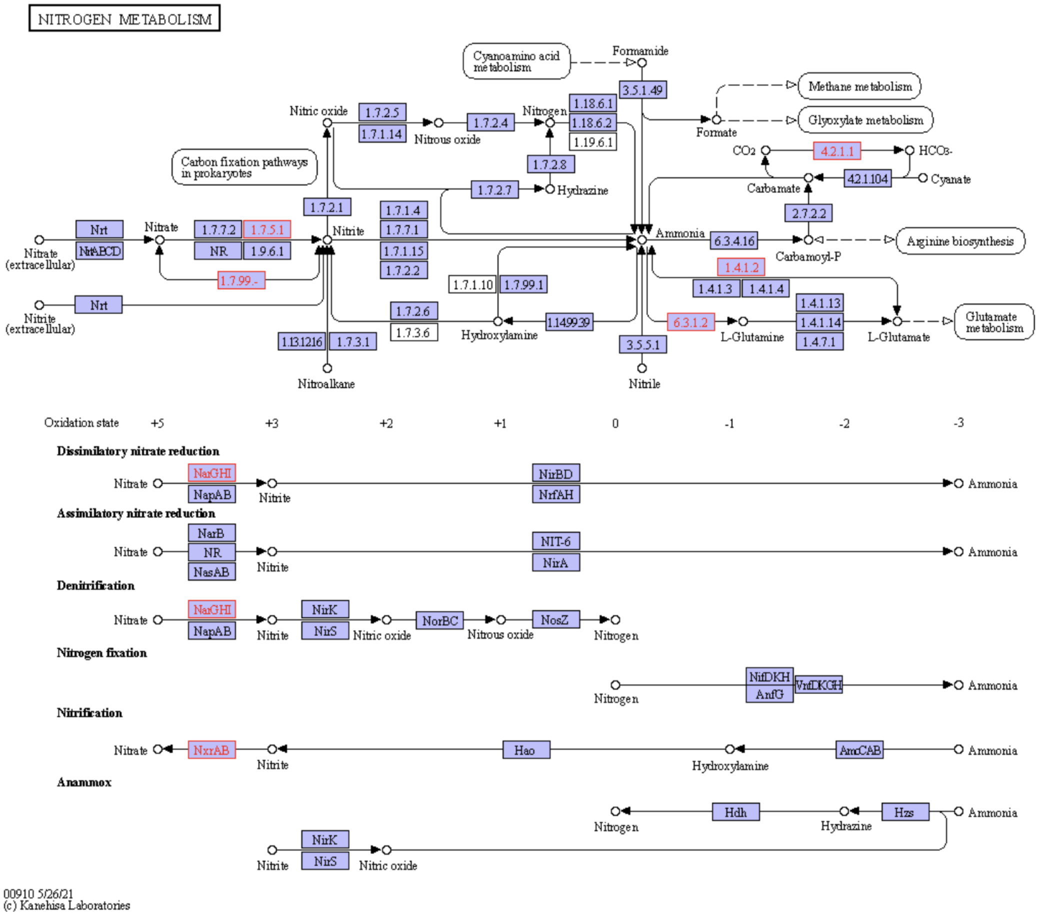


*Figure S8 - Nitrogen metabolism pathway with genes found in drinking water viral population genomes in pink.*

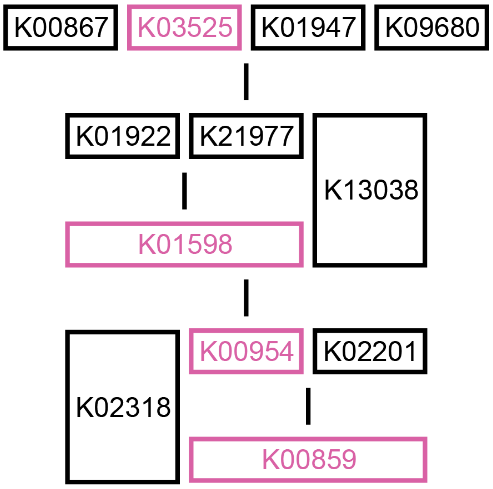


*Figure S9 - Coenzyme A biosynthesis pathway with genes found in drinking water viral population genomes in pink.*
